# Spatial variation in population genomic responses to over a century of anthropogenic change within a tidal marsh songbird

**DOI:** 10.1101/2022.06.10.495648

**Authors:** Phred M. Benham, Jennifer Walsh, Rauri C. K. Bowie

**Affiliations:** Museum of Vertebrate Zoology & Department of Integrative Biology, University of California, Berkeley, Berkeley, CA; Fuller Evolutionary Biology Program, Cornell Lab of Ornithology, Cornell University, Ithaca, NY

**Keywords:** conservation genomics, tidal marshes, Passerellidae, anthropogenic change, museum collections, exome capture

## Abstract

Combating the current biodiversity crisis requires the accurate documentation of population responses to human-induced ecological change. To this end, museum collections preserve a record of population responses to anthropogenic change that can provide critical baseline data on patterns of genetic diversity, connectivity, and population structure. We leveraged spatially-replicated time series of specimens to document population genomic responses to the destruction of nearly 90% of coastal habitats occupied by the Savannah sparrow (*Passerculus sandwichensis*) in California. Spatial-temporal analyses of genetic diversity from 219 sparrows collected between 1889-2017 showed that the amount of habitat lost was not predictive of genetic diversity loss. Despite experiencing the greatest levels of habitat loss, we found that genetic diversity in the San Francisco Bay Area remained relatively high. Over the past century, immigration into the Bay Area from interior populations has also increased. This may have minimized genetic diversity declines, but likely led to the erosion of divergence at loci associated with tidal marsh adaptation. Tracing the genomic trajectories of multiple populations over time provided unique insights into how shifting patterns of gene flow through time in response to human-induced habitat loss may contribute to negative fitness consequences.

## INTRODUCTION

Habitat loss is a primary driver of biodiversity loss [1]. At the genetic level, habitat-induced diversity loss can exacerbate extinction risk through increased mutational load and a reduced capacity to adapt to future environmental change [2-3]. Consequently, the preservation of genetic diversity is now recognized as a major priority for the conservation of global diversity [4]. A predictable relationship between habitat and genetic diversity loss exists that can inform how much of a species’ range must be protected to maintain target levels of genetic diversity [5]. However, across a species’ distribution local adaptation, population size, and gene flow will influence the rate and kinds (e.g. neutral vs. locally adapted variants) of genetic diversity being lost in response to habitat loss and degradation. For instance, habitat deterioration can turn populations into sinks with increased immigration from a source population needed to sustain the population. Changes in rate of gene flow could lead to the evolutionary rescue of a small population [6] or could lead to outbreeding depression if immigrants from source populations are maladapted [7]. Despite mixed theoretical and empirical support for these different outcomes, a detailed understanding of how interactions among different ecological and demographic processes shape patterns of genetic variation in species threatened by habitat loss is lacking. Addressing this gap will be critical for guiding habitat protection and restoration efforts to conserve target levels of genetic diversity.

In this case, looking to past population genetic responses to habitat loss will provide critical insights into the impacts of habitat degradation on genetic diversity. Natural history collections typically preserve specimens collected from the past 150-200 years, a period marked by dramatic human-induced landscape transformation, that can be used to reconstruct population genomic responses to past habitat loss [8-11]. In particular, with the increased ease of generating genomic-scale data from historic DNA sources, analyzing replicated time-series of specimens from across a species distribution will provide important case-studies documenting the role of habitat deterioration in disrupting migration, local adaptation, and drift among populations. To assess how human-induced changes in these meta-population dynamics may impact temporal patterns of genetic diversity, we leverage a spatially replicated time-series of Savannah sparrow (*Passerculus sandwichensis*) specimens that were densely sampled from throughout the state of California over the past 128 years.

The Savannah sparrow is a widespread North American songbird, with 17 subspecies breeding across a range of open habitats from Alaska to Guatemala [12]. Within California, four subspecies span a landscape that has experienced dramatic, but spatially variable, transformations due to human activity (Fig. 1a). The coastal subspecies *P. s. alaudinus* and *P. s. beldingi* primarily occupy tidal marsh habitats in northern and southern California, respectively. These coastal populations exhibit local adaptation to high salinity and flooding in tidal marshes with prior work documenting physiological, behavioral, and genomic divergence from other inland, freshwater-associated California populations of the species [13-17]. Coastal populations have experienced nearly 90% human-induced habitat loss in certain estuaries [18-19] and are now listed as either state endangered species (*P. s. beldingi;* [20]) or bird species of special concern (*P. s. alaudinus;* [21]). In contrast, *P. s. nevadensis* and *P. s. brooksi* of northern and eastern California are of least conservation concern with habitats remaining relatively intact.

**Figure 1:**
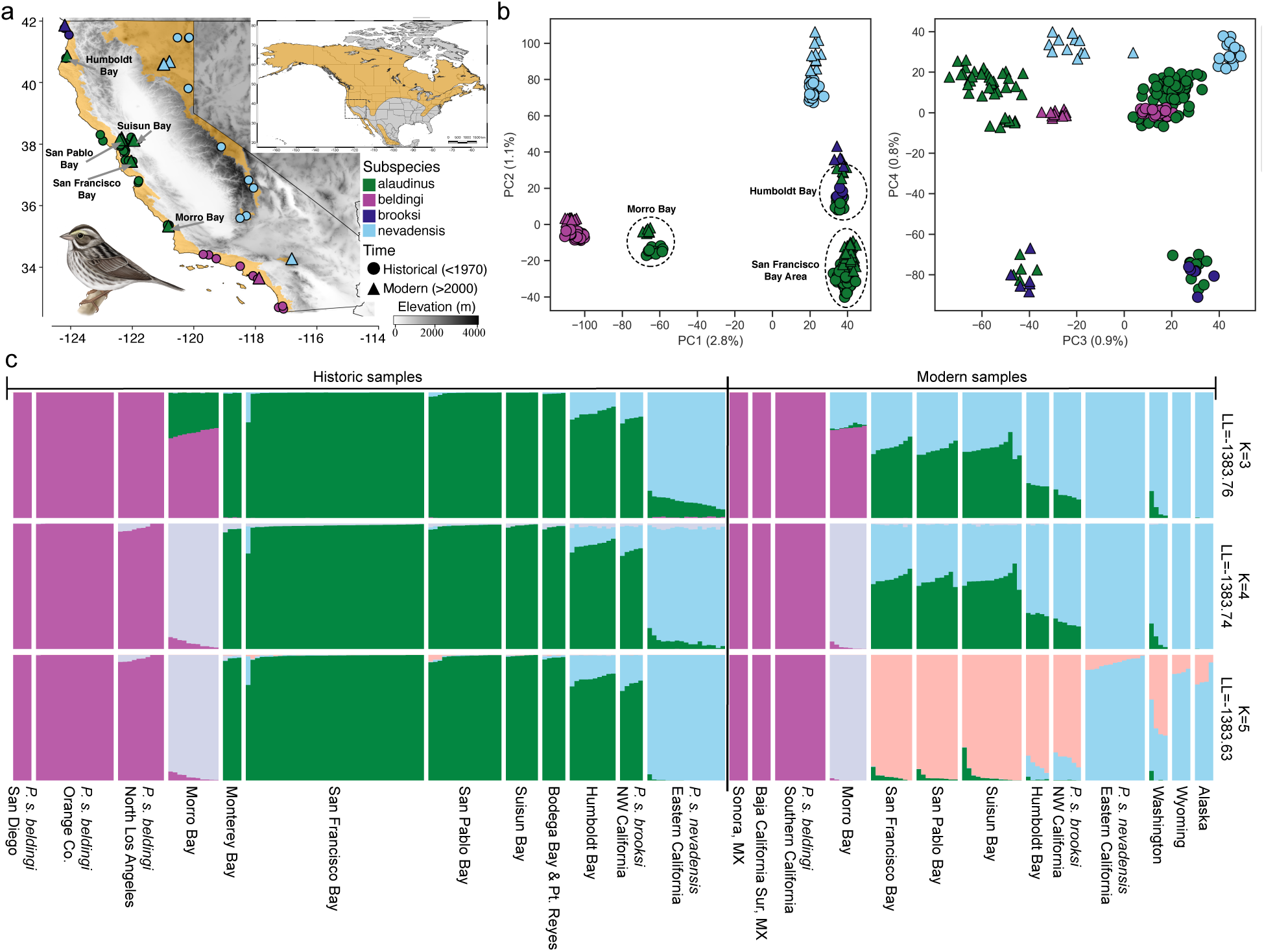
Spatial and temporal patterns of population structure within California populations of the Savannah sparrow (*Passerculus sandwichensis*). **(a)** California breeding distribution of the Savannah sparrow (ochre). Circles represent historical (pre-1970) sampling localities of specimens sequenced in the study, and triangles represent modern (post-2000) sampling localities. Inset shows the extent of the entire North American breeding distribution of the Savannah sparrow in ochre. **(b)** Principal components analysis of the full exome capture dataset (60,765 SNPs). Left panel shows PC1 (2.8% variance explained) versus PC2 (1.1% variance explained) and right panel shows PC3 (0.9% variance explained) versus PC4 (0.8% variance explained). Colors correspond to different subspecies as illustrated in Fig. 1a. Dashed circles denote different clusters corresponding to different populations within *P. s. alaudinus.* **(c)** Results from DyStruct analysis. Runs with clusters specified as K=3-5 all exhibited very similar log-likelihoods and are illustrated here. Colors correspond to different K clusters. Artwork of *P. s. beldingi* individual courtesy of Jillian Nichol Ditner

The contrasting histories of habitat loss experienced by different sparrow populations provide a natural experiment for exploring the temporal dynamics of genetic diversity change in response to human activities. We sequenced genomic data from 219 individuals sampled from 1889-2017, representing nine spatially replicated time-series and all four subspecies. With this dataset, we asked: (1) how has habitat loss impacted patterns of population structure and gene flow among populations? (2) Does habitat loss predict genetic diversity loss? And (3) how have temporal changes in genetic diversity and gene flow among populations impacted loci associated with local adaptation to different habitats? Addressing these questions in concert with time-series data provides a novel framework for evaluating how human-induced ecological change can disrupt historic patterns of local adaptation, gene flow, and population size.

## RESULTS

### Spatial and temporal inference of population structure

We sequenced 145 historical (sampled pre-1970) and 74 modern (post-2000) Savannah sparrow specimens using an exome capture approach (see supplemental results, Fig. S1-S3, and Table S1 for capture performance). Although whole genome sequencing of degraded DNA is increasingly feasible [22-23], target capture approaches have proven to be a highly targeted and cost-effective tool for analyzing temporal genetic change [24-25]. Principial components analysis of all 219 individuals (60,765 SNPs) confirmed that modern and historical samples collected from the same locality clustered together (Fig. 1b). The first principal component explained 2.8% of the variance in the data and separated out three clusters: (1) *Passerculus sandwichensis beldingi* subspecies from south coastal California; (2) birds from Morro Bay in San Luis Obispo County (*P. s. alaudinus*); and (3) birds from the rest of California (*P. s. alaudinus, brooksi,* and *nevadensis*). The second principal component explained 1.1% of the variance in the data and divided birds from the *P. s. nevadensis* subspecies in eastern California, from birds in northwest California (*P. s. alaudinus* and *P. s. brooksi*), and the remaining birds in the *P. s. alaudinus* subspecies from the Bay Area and central California coast. The third principal component (0.9% of variance) separated modern from historical samples and the fourth component (0.8% of variance) split northwest California birds from other populations. PCA of other datasets (downsampled coverage, CpG sites removed, C->T and G->A sites removed) revealed similar patterns of population clustering (supplemental Figure S4).

To further test for temporal changes in population structure and admixture among populations, we analyzed a dataset of 65,583 unlinked SNPs and 239 individuals (an additional 20 from outside California) within the program DyStruct [26]. DyStruct resembles other model-based clustering programs, but explicitly accounts for the temporal dynamics of allele frequency change due to genetic drift in time-series data. The highest likelihoods were for K=3 to 5 (log-likelihood= -1383.63 to -1383.76; Fig. 1c). All three values of K identified birds from south coastal California and Mexico as a distinct cluster, sparrows from the northern California coast as a second genetic cluster, and birds of eastern California, Wyoming, Alaska, and Washington state forming a third cluster. The main difference between K=3 and 4 was that Morro Bay birds were recognized as a distinct cluster, and at K=5 modern birds from the northern California coast were identified as a distinct cluster. This analysis also revealed a key finding: temporal variation in population structure, with modern birds from the central and northern California coast exhibiting a 17% (Morro Bay) to 64% (Humboldt Bay) increase in the proportion of ancestry shared with inland eastern California birds. These temporal changes point to a recent increase in the level of gene flow from inland eastern California populations into coastal populations.

### Spatial and temporal patterns of genetic diversity

We next explored how variation in the scale of human-induced habitat loss influenced patterns of genetic diversity across sparrow populations. First, effective diversity surfaces estimated in EEMS [27] show similar patterns of genetic diversity among populations in both the historical (<1940) and modern (>2000) datasets (Fig. 2a & 2b) with southern California birds consistently exhibiting lower effective diversity rates across time relative to northern California birds. Second, the southern California birds exhibited significantly lower levels of nucleotide diversity (π) and Watterson’s theta (*θ*) compared to northern birds (Fig. 2c; historic one-way ANOVA analysis [π: F=58.2, df=7, p<<0.0001; *θ*: F=252.9, df=7, p<<0.0001]; modern one-way ANOVA analysis [π: F=101.9, df=7, p<<0.0001; *θ*: F=301.7, df=7, p<<0.0001]).

**Figure 2:**
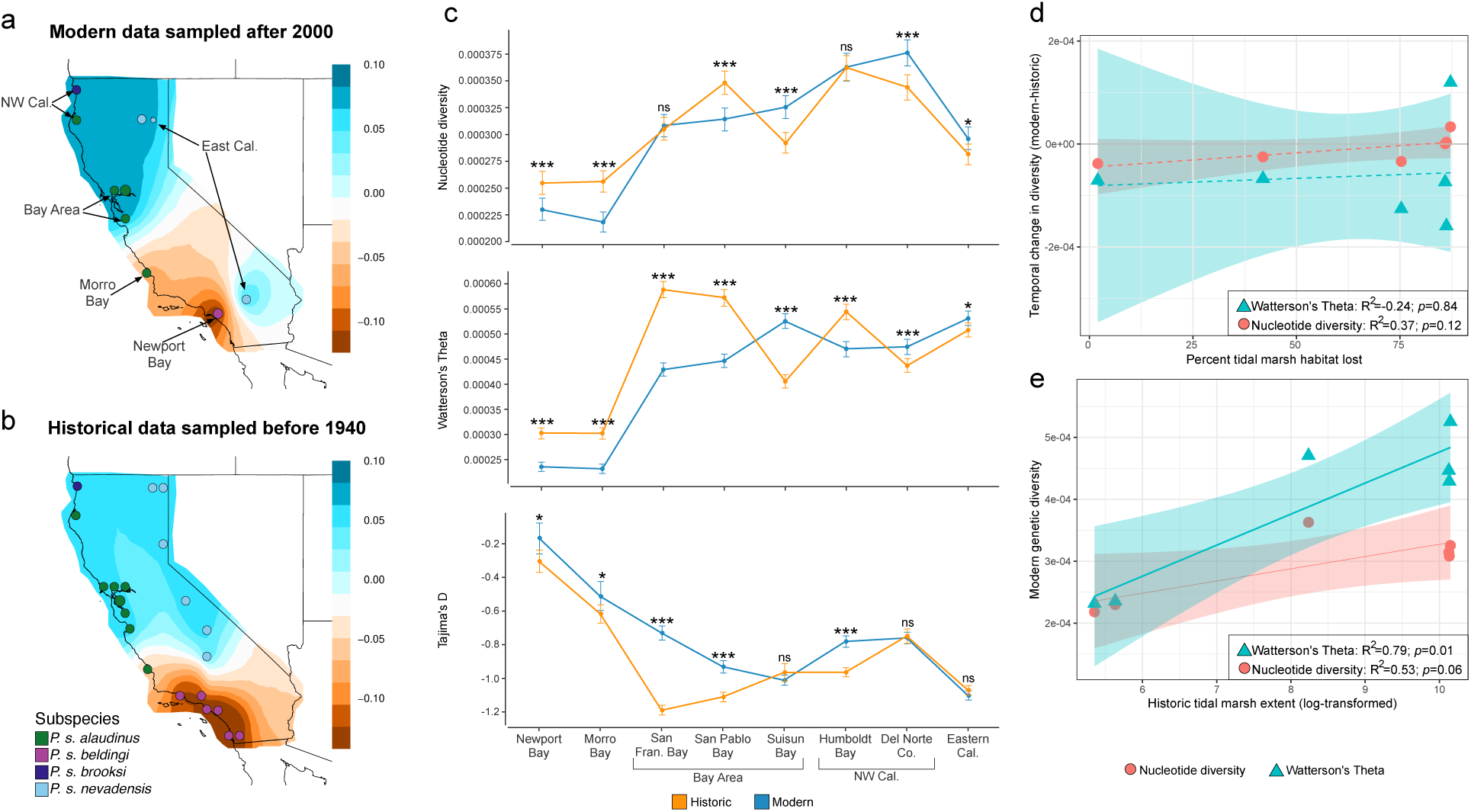
Spatial and temporal patterns of genetic diversity within California Savannah sparrows. **(a)** Modern variation in genetic diversity across California estimated using the Estimated Effective Migration Surfaces (EEMS) program. Contour plots vary from low (dark brown) to high (dark blue) posterior mean diversity rates. **(b)** Historic variation in genetic diversity across California estimated using EEMS. **(c)** Plots comparing historical (orange) and modern (blue) estimates of nucleotide diversity, Watterson’s theta, and Tajima’s D across all populations. For both **b** & **c** plots are based on a downsampled dataset with C->T and G->A mutations removed. Asterisks in **b** & **c** denote p-values based on t-tests between time groups (ns: non-significant; *p<0.05; **p<0.01; ***: p<0.0001). **(d & e)** Relationship between the amount of tidal marsh habitat lost or available in each of six estuaries and population genetic statistics. Values on y-axis have all been scaled and centered to visualize on same axis. **(d)** The amount of tidal marsh lost (%) over past century is not significantly related to temporal changes in nucleotide diversity (red circle) or Watterson’s theta (blue square) (**e)** Historically, the total amount of tidal marsh habitat present in each estuary significantly predicts modern levels of Watterson’s theta, while nucleotide diversity is marginally non-significant. Contemporary levels of tidal marsh habitat did not predict levels of genetic diversity (Supplemental Table S2).

California’s coastal ecosystems experienced more dramatic habitat loss over the past century [18-19; 21] relative to localities in eastern and northern California. Consistent with these differences in landscape change, we found significant increases in π and *θ* for eastern and northern California populations, but evidence for significant declines in genetic diversity for five out of six tidal marsh populations in coastal California (Fig. 2c). Suisun Bay in the Bay-Delta region of central California was the only coastal population that exhibited significant increases in genetic diversity. Declines in genetic diversity were more evident in *θ* as opposed to π, which contributed to a significant increase in Tajima’s D for all tidal marsh populations (except Suisun Bay). This pattern is indicative of a population undergoing a contraction where rarer variants (as measured by *θ*) are eliminated before overall declines in heterozygosity (measured by π). Estimates of these parameters using datasets that were less conservatively filtered showed similar patterns (supplemental Fig. S5). For Newport Bay and San Francisco Bay trends in genetic diversity metrics were also evaluated across samples taken from three and six time points, respectively. For Newport Bay birds, significant decreases in π and *θ* only occurred between the 1960s and 2010s (supplemental Fig. S6). San Francisco Bay on the other hand showed a long-term increase in Tajima’s D from the early 1900s to the present, driven by decreasing *θ* through time (supplemental Fig. S7).

We also noted significant differences in the magnitude of genetic diversity decline among tidal marsh populations (one-way ANOVA analysis; π: F=294.7, df=5, p<<0.0001; *θ*: F=61.15, df=5, p<<0.0001). To determine whether differences in the amount of habitat loss experienced by different tidal marsh populations explained this variation in genetic diversity loss, we compared estimates of percent marsh habitat lost (data from [19]) for each estuary with temporal change in genetic diversity. The amount of tidal marsh loss varied from 2-87%, but this variation did not significantly explain temporal change in π or *θ* based on simple linear regression models (Fig. 2d). We further tested whether total amount of tidal marsh area (log-transformed) in historic or modern times predicted change in genetic diversity across sites, or whether contemporary extent of tidal marsh explained variation in genetic diversity. All of these relationships were non-significant (Supplemental Table S2). The only significant predictor of modern levels of genetic diversity was the historic area of tidal marsh, which significantly predicted variation in contemporary *θ* (R^2^=0.79; p=0.01) and the relationship with π was nearly significant (R^2^=0.53; p=0.06; Fig. 2e). These results indicate that despite widespread habitat loss across all coastal marshes, the populations from the historically largest marshes continue to maintain the highest levels of genetic diversity. This is true even though populations from some of the most expansive marshes (e.g., San Francisco Bay) experienced the greatest habitat loss and decline in *θ*, but even in modern times maintain levels of *θ* comparable to other northern California populations.

### Temporal change in demography

To quantify the extent of temporal change in genetic diversity and migration rates among populations we next fit a series of demographic models to the three-dimensional site frequency spectrum of eastern California (*P. s. nevadensis*), Bay Area (*P. s. alaudinus*), and Newport Bay (*P. s. beldingi*) birds in GADMA2 [28]. We ran 20 independent runs of GADMA2 for both a historic and modern dataset. For historic birds, the best-fit model (Fig. 3a; Table1; Log-likelihood -1391.83) showed an initial divergence time between Newport Bay and the northern California populations ∼254 kya, followed by divergence between the Bay Area and Eastern California populations ∼32 kya. Following the initial split, birds from southern California (Newport Bay) grew from an effective population size (Ne) of 0.2 to 16,749 before contracting again to 6,675. In contrast, the northern California population remained large with an Ne of 297,826 before splitting into an eastern California population that maintained a constant Ne of 68,438 and a Bay Area population that declined to an Ne of 16,755. Migration rates were roughly symmetrical from eastern California to the Bay Area (1.6e-04) and vice versa (1.3e-04), which translates to a rate of 8.9 migrants per generation (Nm) from the Bay Area to eastern California and Nm=2.7 in the reverse direction. Overall, the top five of 20 models fit to the historic dataset show a similar demographic history and parameter estimates (Table 1).

**Figure 3:**
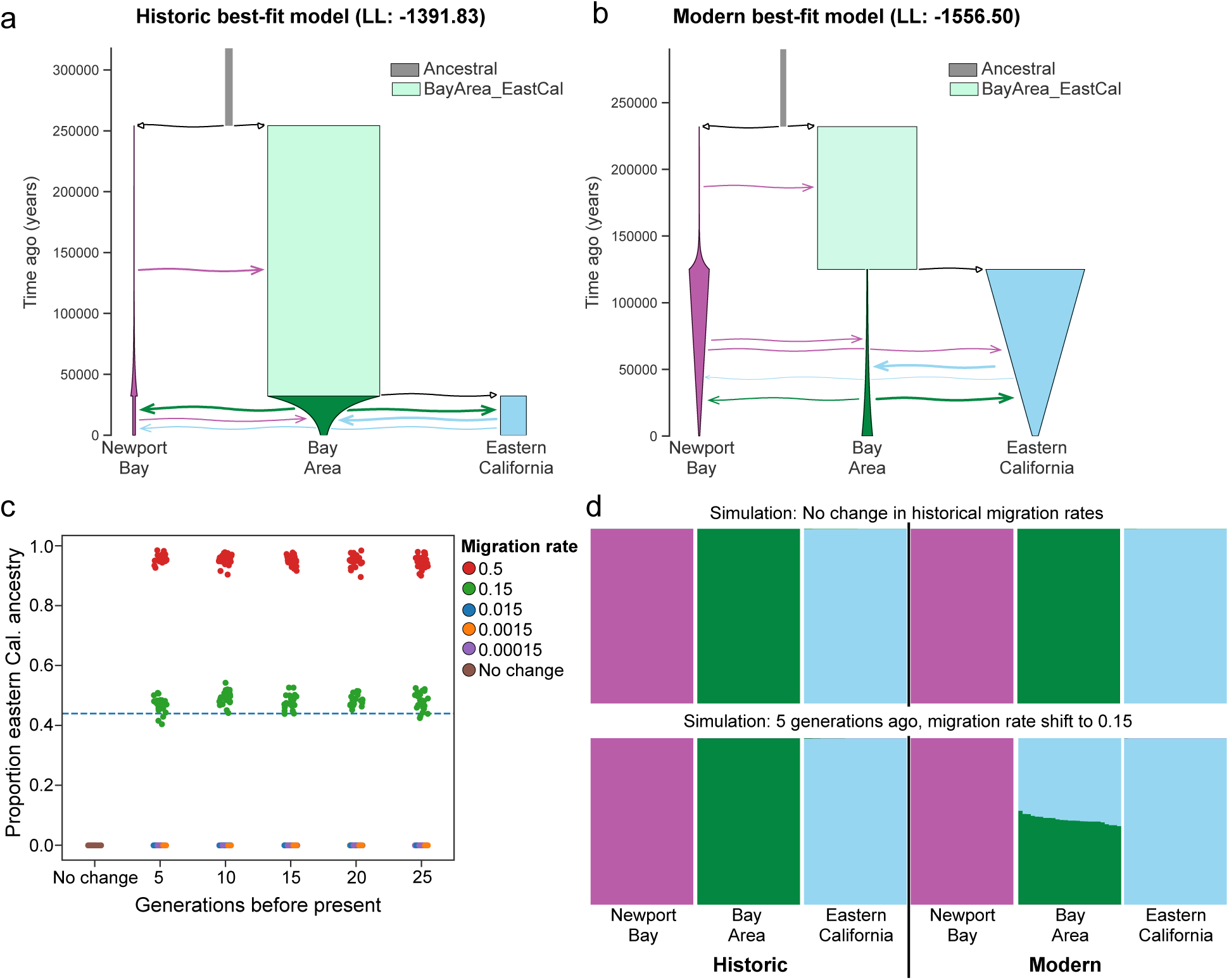
Temporal change in migration rates within California Savannah sparrows. (a) Historic best-fit model estimated with GADMA2. Y-axis shows time of demographic events, width of bars represents effective population size for each population, and arrows signify migration rates from source to destination population forward in time. Arrows are colored by source population and represent continuous migration. (b) Representation of best-fit model to the modern dataset in GADMA2. (c) Coalescent simulation results where the historic best-fit model (Fig. 3a) was used as the baseline demographic history with migration rates changing at varying generations before the present (x-axis) to different rates (colors in legend) from eastern California to the Bay Area. Simulated datasets were analyzed within Dystruct and the proportion of eastern California ancestry in the modern Bay Area population was estimated (y-axis). Dashed blue line represents the average observed eastern California ancestry found in the modern Bay Area population (see Fig. 1c). Migration rates <0.15 did not result in any changes to proportion of eastern California ancestry. (d) Barplots showing ancestry proportions estimated for simulated datasets in Dystruct. Top plot shows the results based on simulation of the historic demographic model (Fig. 3a) with no change in migration rate. Bottom plot shows results from simulated dataset where migration rate shifted 5 generations ago from a proportion of 0.00015 immigrants per generation to a proportion of 0.15 immigrants per generation from eastern California into the Bay Area population.

**Table 1:** Parameter estimates for the top five best-fit models for the historic and modern datasets. Demographic models and parameters were optimized in GADMA2. We report parameters for only the most recent time slice. Models are ordered by likelihood from left to right. For both time periods three populations were measured Newport Bay (n), Bay Area (b), and East Cal (e).

|  | **Historic** |  |  |  |  |
| --- | --- | --- | --- | --- | --- |
| LL | -1391.83 | -1393.98 | -1397.1 | -1398.1 | -1405.83 |
| Tsp1 (years) | 254,219 | 238,351 | 192,733 | 243,921 | 3525,73 |
| Tsp2 (years) | 32,164 | 48,431 | 69,673 | 51,533 | 18,578 |
| NewportBay Ne | 6,675 | 9,341 | 13,180 | 8,878 | 6,618 |
| BayArea Ne | 16,755 | 38,319 | 23,688 | 35,013 | 30,276 |
| EastCal Ne | 68,438 | 84,073 | 102,343 | 77,387 | 54,259 |
| n->b | 1.2E-05 | 2.6E-08 | 2.6E-05 | 3.3E-05 | 0 |
| b->n | 1.1E-04 | 8.5E-05 | 4.4E-05 | 2.8E-08 | 1.2E-04 |
| n->e | 0 | 7.5E-06 | 1.4E-05 | 0.0E+00 | 0 |
| e->n | 1.2E-05 | 0 | 2.7E-06 | 7.2E-05 | 0 |
| e->b | 1.6E-04 | 5.9E-05 | 9.8E-05 | 1.1E-04 | 1.1E-04 |
| b->e | 1.3E-04 | 1.1E-04 | 1.1E-04 | 1.1E-04 | 1.5E-04 |
| b->e mig per gen | 8.90 | 9.25 | 11.26 | 8.51 | 8.14 |
| e->b mig per gen | 2.68 | 2.28 | 2.32 | 3.85 | 3.33 |
|  | **Modern** |  |  |  |  |
| LL | -1556.5 | -1583.13 | -1585.11 | -1585.43 | -1595.09 |
| Tsp1 (years) | 232,053 | 276,701 | 310,210 | 313,395 | 343,840 |
| Tsp2 (years) | 125,040 | 31,727 | 5,803 | 27,926 | 3,694 |
| NewportBay Ne | 4,923 | 3,091 | 3,290 | 3,149 | 2,379 |
| BayArea Ne | 57,309 | 41,061 | 37,157 | 426,253 | 54,707 |
| EastCal Ne | 31,638 | 212,932 | 37,029 | 17,969 | 34,174 |
| n->b | 4.2E-05 | 1.5E-05 | 5.2E-05 | 2.6E-05 | 7.2E-05 |
| b->n | 4.5E-05 | 8.4E-05 | 1.0E-04 | 6.1E-05 | 1.2E-04 |
| n->e | 2.6E-05 | 1.9E-20 | 0 | 5.9E-06 | 0 |
| e->n | 6.8E-09 | 2.6E-05 | 1.9E-05 | 7.7E-05 | 0 |
| e->b | 1.0E-04 | 1.0E-04 | 2.4E-08 | 1.5E-04 | 0 |
| b->e | 1.1E-04 | 4.0E-05 | 0 | 3.1E-05 | 0 |
| b->e mig per gen | 3.48 | 8.52 | 0.00 | 0.56 | 0.00 |
| e->b mig per gen | 5.73 | 4.11 | 0.0009 | 63.94 | 0.00 |

The best-fit model for the modern dataset (Fig. 3a; Table1; Log-likelihood -1556.50) showed a similar initial divergence time of ∼232 kya, but a much earlier divergence time between eastern California and the Bay Area (∼125 kya). Newport Bay birds showed a similar history of expansion and decline to a smaller Ne of 4,923. In contrast to the historic dataset, eastern California showed a smaller Ne (31,638) than Bay Area birds (57,309). This difference was consistent across 4 of the 5 top models estimated for the modern dataset. Migration rates were also inferred to be symmetrical between eastern California and the bay area (1.0e-04 and 1.1e-04) with a slightly greater Nm per generation (5.73) from eastern California to the Bay Area than in the opposite direction (Nm=3.48), a reversal from the historic dataset. Unlike the historic dataset, the top 5 models showed extensive variation in a number of parameter estimates, notably related to divergence time between eastern California and the Bay Area (range 3.6 kya to 125 kya), Bay Area Ne (37,157 to 426,253), and Nm from eastern California to the Bay Area (0 to 63.94).

The reasons for the increased variation among models in the modern dataset was unclear, however, greater variation could relate to recent and unaccounted for changes in demographic history due to population declines or shifts in migration rate that were not captured by the demographic model. The latter possibility was previously suggested by DyStruct analysis where a significant increase in eastern California ancestry was observed in modern day Bay Area populations (Fig. 1c). To test this, we performed a series of coalescent simulations in msprime 1.2.0 [29] parameterized by the best-fit historic demographic model. Analysis of 26 simulated SNP datasets in DyStruct showed that a shift to a migration rate of 0.15 from eastern California to the Bay Area 5-25 generations ago would be necessary to match observed proportions of eastern California ancestry in Bay Area birds (Fig. 3cd). Simulation results provide strong support for observed DyStruct patterns being the result of significant changes in the rate of gene flow from eastern California to the Bay Area as opposed to other processes such as genetic drift.

### The influence of demographic change on locally adapted loci

Finally, we aimed to understand how the inferred changes in gene flow between tidal marsh and freshwater adapted (eastern California) populations of the Savannah sparrow may have impacted divergence in loci putatively associated with local adaptation to tidal marsh habitats. We first compared genome wide mean Fst divergence from eastern California in the four tidal marsh populations (Humboldt Bay, Bay Area, Morro Bay, Newport Bay) for historic and modern datasets. Modern values of Fst divergence showed increases in Morro Bay from 0.054 to 0.062, but a sharp decrease in mean Fst at Humboldt Bay from 0.014 to -0.002 and more modest changes in the other two populations (Table 2). We next estimated Fst and Dxy in sliding windows to identify loci exhibiting evidence for divergent selection between eastern California populations and each of the four tidal marsh populations in the historic dataset. Additionally, we identified SNPs that varied significantly in association with salinity variation across populations using latent factor mixed models (LFMM [30]). Across the four populations, we identified 67 outlier windows in the historic dataset that were in the 95th percentile of Fst, Dxy, and LFMM analyses (Fig. 4a).

**Table 2:** Mean Fst divergence between eastern California and each of four tidal marsh populations of the Savannah sparrow in both the historic and modern datasets.

| **Population** | **historic**<br>**sample size** | **modern**<br>**sample size** | **Number**<br>**SNPs** | **Historic**<br>**mean Fst (sd)** | **Modern**<br>**mean Fst (sd)** |
| --- | --- | --- | --- | --- | --- |
| Newport Bay | 10 | 11 | 4495 | 0.098 (0.008) | 0.099 (0.009) |
| Morro Bay | 11 | 8 | 5408 | 0.054 (0.003) | 0.062 (0.006) |
| Bay Area | 31 | 37 | 12021 | 0.015 (0.001) | 0.013 (0.001) |
| Humboldt Bay | 10 | 5 | 6466 | 0.014 (0.001) | -0.002 (0.002) |

**Figure 4:**
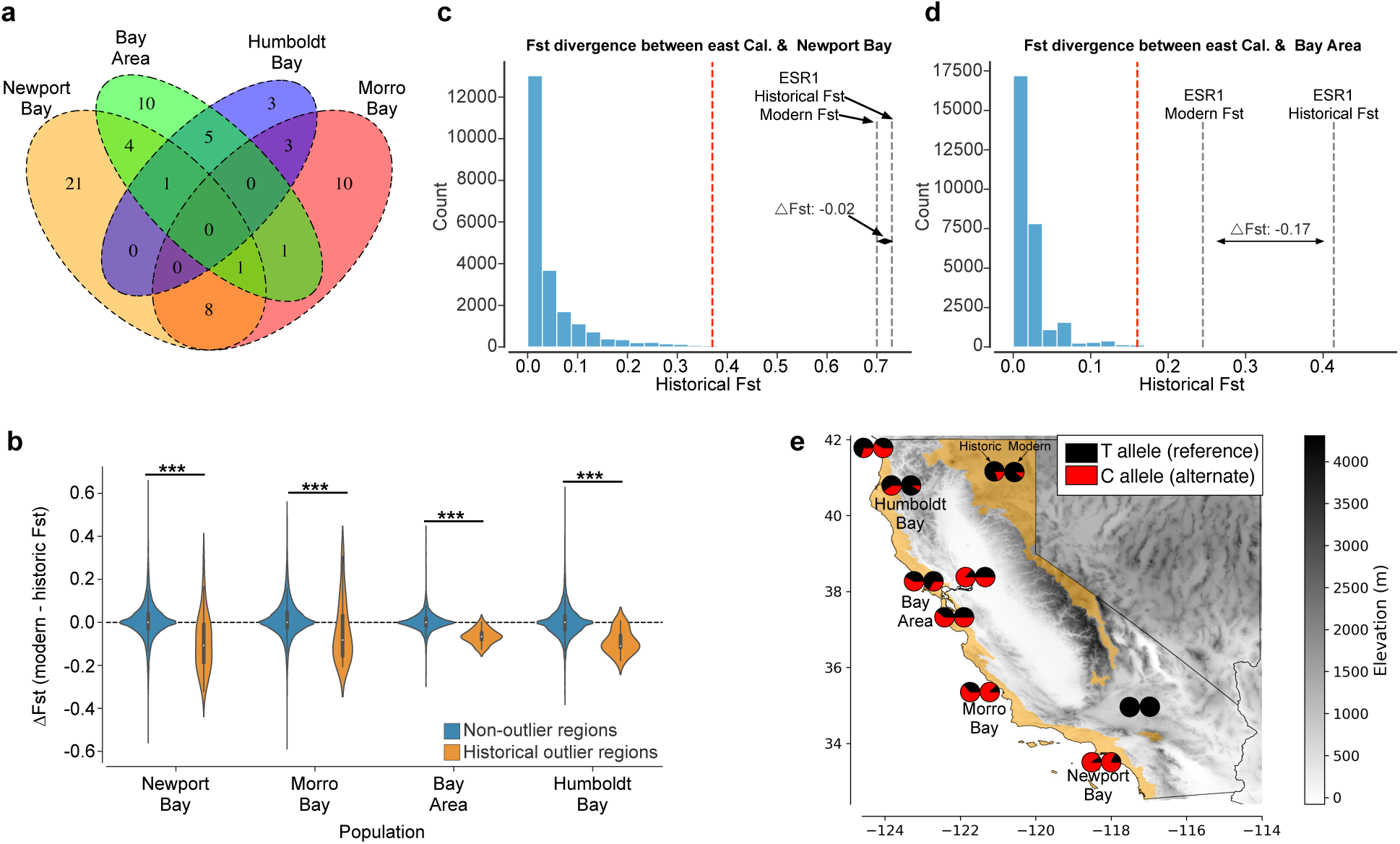
Temporal variation in Fst divergence estimated for genes found to be outliers in Fst, Dxy, and LFMM analyses. **(a)** Venn diagram showing overlap in genes with outlier SNPS across the four tidal marsh populations. **(b)** Change in Fst divergence between eastern California and four tidal marsh populations of Savannah sparrow. ΔFst for each locus was estimated as the difference between historic and modern Fst divergence with positive values indicating increasing Fst divergence through time. Asterisks denote p-values from t-tests of ΔFst between outlier and non-outlier SNPs (ns: non-significant; *p<0.05; **p<0.01; ***: p<0.0001). **(c-d)** Distribution of per SNP Fst estimates for historical comparison between Newport Bay (*P. s. beldingi*) and Bay Area (*P. s. alaudinus*) with eastern California (*P. s. nevadensis*). Red dashed line denotes the mean 99th percentile threshold for calling outlier SNPs. Gray dashed lines represent modern and historic Fst estimates for a C->T mutation in the 3’ UTR region of the final exon of the Estrogen Receptor 1 gene (ESR1). **(e)** Spatial variation in allele frequencies for the ESR1 T->C SNP across nine sampling localities with paired modern and historic sampling. Pie charts on left illustrate historical allele frequencies and pie charts on the right modern allele frequencies.

To assess whether these outlier loci experienced more or less change in Fst relative to non-outlier loci we compared patterns of ΔFst among tidal marsh populations. ΔFst measures the difference in Fst divergence from eastern California between the modern and historic datasets with negative values signifying decreasing divergence through time. This analysis revealed that all four tidal marsh population have experienced significantly greater declines in Fst divergence within outlier loci relative to non-outlier loci (Fig. 4b). Observed patterns of ΔFst in the Bay Area and Newport Bay significantly exceeded the declines predicted by simulations of a neutral demographic history with no demographic changes over the past 100 years (Supplemental Fig. S8); however, ΔFst for Bay Area outlier loci remained significantly greater than ΔFst estimated from a simulated history of recent migration rate change (Supplemental Fig. S8).

No genes were found to be outliers in all four coastal populations, but two genes were outliers in three populations (Fig. 4a; supplemental Table S2). This includes the estrogen receptor 1 gene (ESR1), which was an outlier in all populations except Humboldt Bay. Moreover, a T->C mutation in the 3’ UTR region in the final exon of ESR1 exhibited the greatest Fst divergence in both Newport Bay (Fig. 4c; Fst=0.73) and the Bay Area (Fig. 4d; Fst=0.41). This SNP exhibited large temporal declines in the Bay Area (ΔFst=-0.17), which was likely due to decreasing frequencies of the derived C allele of this SNP in the Bay Area (change in frequency -0.24; Fig. 4e) as opposed to allele frequency change in the ancestral T allele that occurs at higher frequency in eastern California. Again patterns at the ESR1 locus are consistent with temporal changes in gene flow eroding divergence at potentially locally adapted loci.

Increased gene flow from inland eastern California populations into northern coastal populations should also lead to the increased homogenization of allele frequencies among populations towards the present. Further, if outlier loci associated with local adaptation to tidal marshes exhibit increased resilience to gene flow then we would expect less homogenization of allele frequencies at these loci relative to non-outlier loci. We tested these predictions by estimating the degree of correlation between historical deviation in allele frequency and change in allele frequency through time for each SNP [31]. In this analysis, stronger correlations will be indicative of greater homogenization due to higher levels of gene flow. For each of the four tidal marsh populations, we estimated deviation from mean allele frequency of the tidal marsh and eastern California populations as 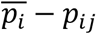. Where 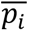 is the mean allele frequency of the i^th^ SNP between inland eastern California and the j^th^ tidal marsh population and *p*_*ij*_ is the allele frequency of the i^th^ SNP in the j^th^ tidal marsh population. Difference in allele frequency change through time was estimated for each SNP in each tidal marsh population as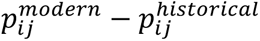. We used permutation tests to test for significant differences from zero (no correlation) as well as between outlier and non-outlier SNPs. This showed that correlations between historic deviation and temporal change in allele frequency were significantly different from zero for both the Bay Area (mean correlation: 0.17; p-value=0.015) and Humboldt Bay tidal marsh populations (mean: 0.30; p-value=0.001), but not significant for Morro Bay (mean: 0.16; p-value=0.08) or Newport Bay (mean: 0.05; p-value: 0.33; Fig. 4a). Second, we found that outlier loci exhibit significantly more positive correlations relative to non-outlier loci across all four coastal populations (Fig. 4a). Finally, the degree of correlation varies significantly with the extent to which inland eastern California ancestry increased in each tidal marsh population through time (r^2^=0.92; p-value=0.03; Fig. 4b). This result remained regardless of the threshold used to identify outlier SNPs (thresholds: 0.9, 0.95, 0.99th percentile of Fst divergence). In sum, ΔFst and temporal allele frequency change results both indicate that the inferred increase in gene flow levels from the freshwater-adapted, eastern California population into tidal marsh populations eroded divergence at loci potentially associated with adaptation to a high salinity environment.

## DISCUSSION

There is a pressing need to understand the ecological and demographic factors that influence population genetic responses to habitat loss and environmental change. Here, we leveraged spatially replicated, time-series of specimens to document the impacts of habitat loss in tidal marsh populations of the Savannah sparrow in California. We found that 5 of 6 tidal marsh populations exhibit evidence for declines in genetic diversity, in line with expectations that coastal populations would experience the greatest overall loss. However, the amount of tidal marsh habitat lost was not linked to patterns of genetic diversity declines (Fig. 2d). Rather, we showed that: (1) historic levels of genetic diversity in southern California were lower than northern California (Fig. 2a-c); (2) modern levels of genetic diversity were correlated with the historic extent of tidal marsh habitat in each estuary (Fig. 2e); and southern populations have likely remained small since divergence from northern populations ∼250 kya (Fig. 3a). Further, despite experiencing nearly 90% tidal marsh habitat loss, populations from the Bay Area (San Francisco, San Pablo and Suisun Bays) still exhibit similar levels of genetic diversity to other northern California populations (Fig. 2c). While habitat loss and fragmentation likely played a primary role in observed declines of genetic diversity, our analyses point to more complex demographic dynamics contributing to overall patterns of changing genetic diversity among these populations.

In other species, a disconnect between habitat loss and genetic diversity declines has been attributed to a range of different demographic, life history, ecological, and behavioral factors [33-34]. Many of these factors are unlikely to contribute to the observed differences in genetic diversity between southern and northern tidal marshes as generation time, body size, and migratory behavior are all similar across coastal populations. Differences in the degree of specialization to tidal marshes could contribute to differences in genetic diversity declines. Tidal marsh habitats are the stronghold for *P. s. alaudinus* populations in northern California, but this subspecies can regularly be found breeding in coastal grasslands [35]. This greater niche breadth may buffer them from tidal marsh habitat losses. Indeed, populations from Humboldt Bay shifted from tidal marsh into pasture habitats as pasture land expanded at the expense of tidal marsh habitat [21].

Reduced genetic diversity declines in the Bay Area may also be due to temporal changes in gene flow rates among populations. Population structure (Fig. 1c), demographic (Fig. 3; Table 1), and simulation analyses (Fig. 3c-d) all support a recent shift to greater immigration from eastern California into the Bay Area. These results support theoretical work that habitat loss and subsequent population declines can leave populations more susceptible to hybridization and gene flow [36]. Further, differences in Ne between populations will influence the direction of gene flow with greater gene flow expected from larger into smaller populations [37]. Our empirical results and theoretical work [38-39] together suggest that habitat loss may have led to declining local recruitment and population growth within the Bay Area, resulting in the Bay Area becoming a sink population, sustained by migration from eastern California.

The impact of gene flow on the persistence of natural populations is hotly debated in the conservation genetics literature [40-42]. Gene flow can be crucial for preventing or reversing inbreeding depression [43-45]. Gene flow can also contribute critical standing variation for adaptation and may be important for future responses to climate change [46]. However, gene flow could also play a role in eliminating divergence between populations and may lead to outbreeding depression [7]. Our data show that coastal populations experiencing higher levels of gene flow do exhibit reduced declines in π (but not Watterson’s *θ*) over time. However, we also show that increased immigration into the Bay Area through time likely contributed to the homogenization of outlier regions associated with tidal marsh adaptation (Fig. 4, 5). Our findings contrast with a recent study on Trinidadian guppies, which found that allele frequency differences in outlier loci were more resistant to experimentally-induced gene flow than non-outlier loci [45]. This study also showed that increasing gene flow from main-stream environments led to increased fitness of headwater guppy populations, but the study was performed only for a limited time frame of ∼6 generations. It will be critical for the future management of the tidal marsh sparrow populations to determine whether increased gene flow contributes to reduced fitness and outbreeding depression. However, the current study remains an important natural replication of the guppy experiment showing that shifting gene flow rates in response to human-induced habitat degradation can lead to the greater erosion of locally adapted loci.

**Figure 5:**
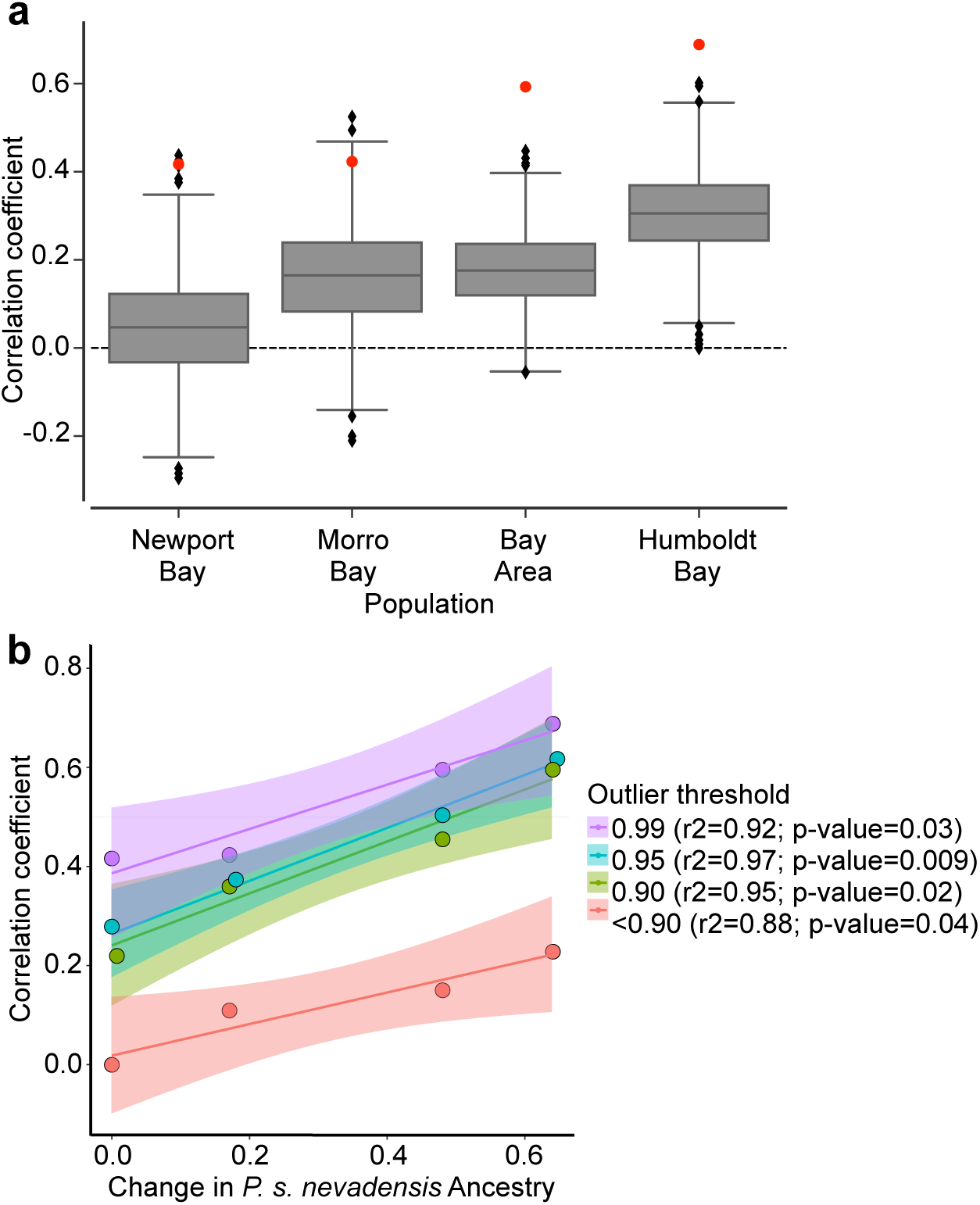
**(a)** Correlations between the historical deviation from mean allele frequency and direction of temporal change in each tidal marsh population. More positive correlations indicate greater changes in allele frequency towards the mean due to higher levels of gene flow. Red circles show observed correlation for outlier SNPs and gray box plots indicate the null distribution from 1000 permutations of non-outlier SNPs. **(b)** Regression analyses showing relationship between change in the proportion of ancestry from *P. s. nevadensis* populations in each tidal marsh population and the correlation coefficient for outlier SNPs shown in (a). Ancestry proportions from DyStruct analysis (Fig. 1c). Different colors represent SNPs inferred to be outliers from different percentile thresholds with red points showing data from non-outlier SNPs.

Linking outlier loci from genome scans to performance and fitness is a persistent challenge in studies of non-model organisms [47]. Despite this caveat, over 40% of identified outlier loci may contribute to increased osmoregulatory performance in high salinity tidal marsh environments via their known roles in kidney development, function, and pathology (Supplemental Table S3). Only the estrogen receptor 1 (ESR1) and IgGFc-binding protein-like (FCGBP-like) genes were found to be outliers across multiple populations. The ESR1 gene in birds has primarily been linked to differences in aggressive behavior [48-49]. However, ESR1 is widely expressed in the mammalian kidney [50] and contributes to the generation of important osmoregulatory phenotypes matching those found in tidal marsh sparrows [16; 51]. These include larger kidneys [52] and the production of a more concentrated urine through the inhibition of Aquaporin-2 expression in the collecting duct of rodent kidneys [53]. Salinity and temperature are expected to increase in northern California in the near future and could exacerbate osmoregulatory stress experienced by these sparrows [54]. If the ESR1 and other candidate genes identified in this study contribute to important osmoregulatory functions, declining allele frequencies due to gene flow may reduce the adaptive capacity of these Bay Area birds.

### Conclusions

Research on geographic variation has long motivated the growth of natural history collections. Consequently, museum collections now provide abundant material for documenting the impact of human-induced ecological change on temporal changes in migration, population size, and selection. Taking full advantage of this unique strength of collections across broad taxonomic scales will be essential for developing a more robust understanding of plant and animal responses to anthropogenic change and translating this knowledge into improved management strategies. Our work demonstrates the power of analyzing these replicate time series of specimens to reconstruct how habitat loss and demography have interacted over the past century to shape population genetic change. Specifically, we provide novel evidence for human-induced habitat loss transforming a locally adapted population from a source population to a sink with associated declines in divergence of locally adapted loci.

## METHODS

### Sample library prep, capture, and sequencing

DNA was extracted from the toepads of 145 historic specimens and blood and tissues from 94 modern samples (74 from California and 20 from outside California; Dataset S1). Historical toepad samples were extracted using a phenol-chloroform protocol in a lab space dedicated to working with degraded DNA. Historic samples were treated with the USER enzyme to mitigate C->T base misincorporation [55]. Modern samples were extracted using a Qiagen DNeasy extraction kit (Valencia, California) and randomly sheared using a qSonica Q800R3 Sonicator (Newtown, CT). For all samples, libraries were prepared for sequencing using a Kapa Illumina Hyper Prep kit (see supplemental methods). DNA from prepared libraries was captured using a custom designed exome capture array (see supplemental methods). For captures, 500 ng of DNA from each library was pooled and hybridized in solution to the SeqCap EZ probes following the manufacturer’s protocols (Roche Sequencing Solutions, Inc, Pleasanton, CA). Captured libraries were sequenced on two lanes of NovaSeq S4 and a single lane of NovaSeq SP at the University of California Berkeley Vincent J. Coates Genomics Sequencing Lab. De-multiplexed reads were filtered and aligned to the reference using the seqCapture pipeline (https://github.com/CGRL-QB3-UCBerkeley/seqCapture/tree/master/scripts). We randomly downsampled the number of reads included in bam files of the modern, higher-coverage samples using the DownsampleSam function in Picard tools (https://broadinstitute.github.io/picard/). Variants from resulting bam files were identified and called using the ’mpileup’ and ’call’ functions in bcftools [56] (see supplemental methods for additional filtering, alignment, and variant calling details).

### Population structure analyses

Principal components analysis of the full, exon, and nongenic datasets resulting from different filtering protocols (supplemental methods) was performed using the python package scikit-allel v. 1.2.0 [57]. For all SNP datasets, we removed singletons and pruned SNPs that were in linkage (r=0.25) in 2500 bp blocks. The estimated effective population size (Ne) in DyStruct was set to 100,000 after preliminary analyses showed that results were robust to varying the Ne setting from the smaller (25,000) to larger population sizes (500,000) previously reported for this species [58]. Generation time was set to 2.2 years [59] and we binned samples into eight 10-year windows spanning 48 generations from 1890 (generation 0) to 2017 (generation 48). We ran DyStruct three times independently for K = 1 to 12, where K is the number of pre- defined population clusters. The most likely value of K was identified using a cross-validation, hold-out method within the program.

### Genetic diversity and demographic analyses

We compared historical and modern patterns of migration and genetic diversity across California with Estimated Effective Migration Surfaces in the program EEMS [27]. For both the modern and historic datasets, input files were generated using the bed2diffs function in EEMS. We ran three independent runs of the MCMC chain. Each chain started from a different random seed and ran for 2 million iterations of burn-in followed by 5 million iterations. A population grid density of 250 demes was selected and we adjusted the qEffctProposalS2 (0.01) and qSeedsProposalS2 (0.2) settings upwards to ensure a proposal acceptance rate between 10-40%. We used the R package rEEMSplots to assess model fit and convergence among the three independent chains.

To match sequence depth between modern and historical samples, population genetic statistics were estimated using a dataset with downsampled coverage in modern samples and the removal of C->T, G->A mutations. We estimated Tajima’s D, nucleotide diversity (π), and Watterson’s Theta (*θ*) for each exon (n=9,971 loci) and non-genic target region (n=1,024 loci) in the dataset using scikit-allel v. 1.2.0 [57]. We also estimated these metrics across three and six time points in Newport Bay (*P. s. beldingi*) and San Francisco Bay (*P. s. alaudinus*), respectively. To estimate the magnitude of temporal change in each metric we subtracted the historic from the modern value. Within each population we assessed whether significant differences existed between sampling points using ANOVA and post-hoc TukeyHSD tests or t-tests depending on the number of time points sampled. All statistical analyses were performed in R version 3.5.1 (https://www.r-project.org/).

We used GADMA2 [28] to identify the demographic model that best fit the three dimensional site frequency spectrum (SFS) of eastern California (*P. s. nevadensis*), Bay Area (*P. s. alaudinus*), and Newport Bay (*P. s. beldingi*) birds. We ran separate analyses for a historic dataset and a modern dataset of these three populations. For separate historical and modern datasets, we generated the SFS from both exon and non-genic data, however, to restrict our analyses to putatively neutral loci we eliminated regions of the genome found to be under selection by our analyses below. The final filtered vcf file was converted to the *Moments* input format using a perl script from (https://github.com/wk8910/bio_tools/blob/master/ 01.dadi/00.convertWithFSC/convert_vcf_to_dadi_input.pl). The final datasets included 17 individuals of *P. s. nevadensis,* 37 *P. s. alaudinus,* and 10 *P. s. beldingi* in the historical dataset, and 13 *P. s. nevadensis,* 31 *P. s. alaudinus, and* 11 *P. s. beldingi* in the modern dataset. Within GADMA2 these datasets were downprojected to include an equal number of samples for each population with 10 for the modern and 11 for the historic population. The final input datasets included 26,080 and 21,758 SNPs for the historical and modern data, respectively.

Within GADMA2, we used *Moments* [60] as an engine for local optimization. We initialized the global search with a simple structured model (1,1,1) that allowed for one demographic event in between divergence times of the three populations. We allowed for asymmetric migration among all populations, and population size changes that were linear, exponential or sudden. For each dataset, we performed 20 independent runs of the global search optimization in GADMA. Since all optimizations produced a model with the same number of parameters top models were identified based on their likelihood. Demographic parameter values were estimated from the value of theta (4*N_e_µL*; where *L* is sequence length) based on a generation time of 2.2 years for the Savannah sparrow [59], an estimated germline mutation rate of 4.6e-9 for another songbird species, *Ficedula albicollis* [61]. The total length of sequence (*L*) from which SNPs were called was estimated as: total sequence length * (SNPs in SFS/ total SNPs pre-filtering). This resulted in a length estimate of 6.5 Mb for the historic dataset and 4.1 Mb for the modern dataset.

### Coalescent simulations

We performed a series of coalescent simulations in msprime 1.2.0 [29] to confirm whether recent changes in migration rates between coastal and interior populations of the Savannah sparrow could explain inferences from DyStruct. We used the historical best-fit GADMA model to parameterize our demographic model in GADMA. In total we performed 25 separate simulations that included all combinations of five different timings of migration rate changes (5,10,15,20,25 generations) and five different migration rates from eastern California to the Bay Area (0.00015, 0.0015,0.015,0.15, 0.5). We ran a final simulation that involved no change in migration rate from the historical best-fit demographic model. For each simulation we sampled 25 individuals from the Bay Area, Newport Bay, and eastern California at 50 generations before the present and another 25 individuals from each of the populations at the present time. This sampling scheme closely resembles the earliest and latest sampling times for DyStruct analyses on observed data. For each of the 26 simulations we ran 60,000 replicates of the demographic model sampling from a sequence length of 650 to approximate the ascertainment scheme of our capture dataset. For each of the 60,000 replicated tree sequences mutations were added using default parameters in msprime and a mutation rate of 4.6e-9 for passerine birds [61]. To generate a genotype matrix from these simulated mutations we randomly selected a single variant from each of the 60,000 replicates that was present in 2 or more individuals. The resulting genotype matrix for each simulation was analyzed in DyStruct to test which parameter combinations most closely approximate observed changes in ancestry proportions. We ran Dystruct for each simulation with k=3 and the pop size parameter set to 70,000.

### Spatial and temporal selection analyses

We identified putative outlier loci using the downsampled dataset with C->T and G->A sites removed, a maf filter of 0.05, and only retained sites with no missing data. We divided the dataset into five populations: (1) freshwater-adapted populations of eastern California (*P. s. nevadensis*); (2) Humboldt Bay estuary (*P. s. alaudinus*); (3) San Francisco Bay estuary (*P. s. alaudinus*); (4) Morro Bay estuary (*P. s. alaudinus*); and (5) Newport Bay estuary (*P. s. beldingi*). Each of these five populations were further divided into historic (sampled 1900-1920) and modern (sampled after 2000) datasets. First, we estimated Hudson’s Fst [62] and Dxy between salt and freshwater-adapted populations for 5Kb windows with a step size of 2.5 Kb following approaches used to analyze other exome datasets [63]. Third, we employed a Latent Factor Mixed Models approach (LFMM; [30]) to identify SNPs with significant associations with mean salinity estimates from a previously published dataset [17]. Loci were designated as outliers if the 5kb region was in the top 5% of Fst divergence, top 5% of Dxy divergence, and contained a SNP with a p-value <0.05 based on LFMM analyses for each of the four populations. We compared temporal changes in Fst divergence for each 5kb window by subtracting historical Fst from modern Fst divergence in each locus to estimate ΔFst. We used t-tests to compare whether outlier versus non-outlier regions exhibited significantly different changes in Fst through time. We performed coalescent simulations in msprime 1.2.0 [29] as above with the historic best-fit GADMA2 model used to parameterize simulations. We simulated a demographic history with no change in the demographic parameters over the past one hundred years and one with an increase in migration rate to 0.15 from eastern California to the Bay Area at 15 generations ago. We simulated a sequence of length 100 Mb and estimated Fst divergence in sliding windows identically to the empirical dataset. Measures of historical Fst divergence, and delta Fst were estimated for both the simulated Bay Area and Newport Bay population. These simulated estimates were compared to observed patterns of Fst divergence using ANOVA and Tukey post-hoc tests in r.

To assess the degree of correlation between historical deviations in mean allele frequency and the direction of allele frequency change through time. We estimated allele frequencies using VCFtools [64]. We classified SNPs as outliers if they exceeded the 99th percentile of Fst divergence. We compared Pearson’s correlation of the outlier SNPs to non-outlier SNPs by performing a permutation test where we randomly sampled an equal number of non-outlier to outlier SNPs and calculated correlations 1000 times. We then compared whether the non-outlier loci were both statistically different from 0 (no correlation) or to the correlation estimated for the outlier loci. We used changes in the proportion of eastern California ancestry found in each of the four coastal populations from the DyStruct analysis to approximate temporal increases in gene flow.

## ACKNOWLEDGEMENTS

We thank the following museums and individuals for providing access to their collections for this study: Museum of Vertebrate Zoology (Carla Cicero); California Academy of Sciences (Maureen Flannery, Jack Dumbacher); San Diego Museum of Natural History (Philip Unitt); Los Angeles County Museum of Natural History (Allison Shultz, Kimball Garrett); American Museum of Natural History (Paul Sweet, Tom Trombone); Field Museum of Natural History (John Bates, Shannon Hackett, Ben Marks); University of Washington Burke Museum (Sharon Burkes); University of Montana Zoological Museum (Angela Hornesby). Personnel at the California Department of Fish and Wildlife, California State Parks, and United States Fish and Wildlife Service assisted with access to federal and state lands. Finally, we thank Elizabeth Beckman, Ammon Corl, Mackenzie Kirchner-Smith, Jackie Childers, Rosa Jimenez, Dan Wait, and Cynthia Wang-Claypool for useful comments on the manuscript. This study was supported by an NSF PRFB fellowship (#1812282) to PMB.

