## supplemental materials for "Spatial variation in population genomic responses to over a century of anthropogenic change within a tidal marsh songbird"

### SUPPLEMENTARY METHODS:

**Capture probe design:** To generate sequence data from both historic and modern specimens, we designed a custom capture array for sparrows (family: Passerellidae) using the white-throated sparrow (*Zonotrichia albicollis*) reference genome version 1.0.1 (Tuttle et al. 2016). We used custom Python scripts to select all exons from the white-throated sparrow (*Zonotrichia albicollis*) genome and add 15 bp of flanking sequence to either end of each exon. Exons were further filtered to only those longer than 180 bp and with GC content between 0.3-0.7% (median GC content = 0.418). We also used RepeatMasker v. 4.0 (Smit, Hubley, Green et al. 2015) to remove exons with excessive repeats. In addition to exons extracted from the *Z. albicollis* genome, we included 314 exons from 111 genes from the zebra finch (*Taeniopygia guttata*) genome that were of particular interest due to their potential role in osmoregulation and the oxidative phosphorylation pathway. Following these steps, we retained a total of 56,230 exons from 13,813 genes. These exons had a mean length of 558 bp (min: 119 bp; max: 14,244 bp) and together totaled 31.6 Mb of the target sequence.

We also included a panel of non-coding regions to assess patterns of neutral divergence. We selected regions of the genome that were at least 100 Kb from any exon to ensure that regions were likely not in linkage with any coding region under selection, given mean LD in birds of ~17Kb (e.g., Balakrishnan & Edwards, 2009; Kawakami et al., 2014). We then randomly selected

1000 bp regions that were each 50 Kb from any other target region, as these putatively neutral regions may face fewer constraints from purifying selection and exhibit higher evolutionary rates. To ensure increased capture efficiency in these regions within Savannah sparrows, we mapped 10 low-coverage Savannah sparrow genomes from Walsh et al. (2019) to these non-coding regions using the program aTram 2.0 (Allen et al. 2018), which iteratively blasts sequence data from bam files to target regions of interest. Finally, we exported a consensus sequence of the low coverage Savannah sparrow genomes that mapped to each non-coding region. This resulted in a final panel of 3,443 non-coding regions with a mean length of 763 bp and a total sequence length of 2,627,883 bp. A single fasta file including all coding and non-coding regions was submitted to Roche Nimblegen to develop probes using the custom SeqCap EZ design.

**Extraction, library prep, and sequencing:**

Historical DNA was extracted from 145 museum specimen toe pads using a phenol-chloroform extraction protocol in a dedicated lab space for dealing with historic DNA. Prior to phenol-chloroform treatment toe pads were minced and digested for ~24 hrs in a solution of 20 ul proteinase K, 20 ul DTT, and 300 ul cell lysis buffer in a rotisserie incubator set at 55ºC. DNA was then added to a 2 ml phase lock tube. The sample was washed twice with 300 ul of phenol-chloroform and twice with 300 ul chloroform. We then performed a bead cleanup using 2x volume of SeraMag beads and two washes of 70% ethanol. The final DNA product was re-suspended in 100 ul of 10 mM Tris. DNA was also extracted from 94 fresh tissue and blood samples using the Qiagen DNeasy extraction kit and following the manufacturer's protocols (Valencia, California).

The sequencing of DNA from historical and ancient samples often results in errors related to the deamination of cytosine nucleotides to uracil, leading to the detection of erroneous C->T substitutions in historic samples (e.g. Briggs et al. 2009). In light of this issue, we treated all historic samples with the USER enzyme (New England Biolabs) in order to cleave uracil nucleotides from historic DNA. We treated approximately 1 ug of DNA with 3 ul of USER enzyme and 10 ul of Cutsmart buffer in a 100 ul reaction that was incubated for 3 hours at 37ºC and then subjected to bead-cleanup prior to library preparation.

Modern samples were randomly sheared using a sonication approach with a qSonica Q800R3 Sonicator (Newtown, CT). All samples were sonicated for a total of 12 minutes with an amplitude of 40% and a pulse of 15 seconds on/15 seconds off. We then performed a double-sided bead cleanup using SPRI low ratio beads with 0.5x right-side selection and 0.65x left-side selection ratios in order to ensure that the DNA fragment distribution was in the 300-500 bp range and to increase capture efficiency. DNA from historic samples was not sonicated given the small fragment size distribution generated by the passage of time.

Using a Kapa Illumina Hyper Prep kit, we prepared libraries of both historical and modern samples for capture and Illumina sequencing. In preparing these libraries, we employed a customized protocol using only a quarter volume, and dual indexes, and we pooled the libraries to include 500 ng of DNA from each library. The pooled libraries were then hybridized in solution to the SeqCap EZ probes, washed, and amplified following the manufacturer's protocols (Roche Sequencing Solutions, Inc, Pleasanton, CA). Captured libraries were sequenced on two lanes of NovaSeq S4 and a single lane of NovaSeq SP at the University of California Berkeley Vincent J. Coates Genomics Sequencing Lab. Separate lanes of NovaSeq S4 included both modern and historic samples. The single lane of NovaSeq SP included only historic samples to increase depth of coverage.

**Filtering, alignment, and variant calling:**

De-multiplexed reads received from the Vincent J. Coates genomics core facility at UC Berkeley were processed using the seqCapture pipeline (https://github.com/CGRL-QB3-UCBerkeley/seqCapture/tree/master/scripts). Briefly, Trimmomatic (Bolger et al. 2011) and Cutadapt (Martin 2011) were used to remove low quality bases (quality score <20) and adapter sequence from the raw reads. Only reads with a minimum length of 50 bp following trimming were retained. We removed PCR and optical duplicates using the hts_SuperDeduper function in HTStream (https://s4hts.github.io/HTStream/); merged overlapping paired end reads were merged using FLASH2 (Magoc & Salzberg 2011); and used Bowtie2 (Langmead & Salzberg 2012) to map reads to the human and *E. coli* genomes and to remove any reads that mapped to these potential contamination sources. We then used the FastQC program (Andrews 2010) to evaluate the overall quality of filtered reads, and mapped the filtered reads to the white-throated sparrow genome using BWA (Li & Durbin 2009). Variants from resulting bam files were identified and called using the 'mpileup' and 'call' functions in bcftools (Li 2011). We also generated a dataset with similar mean coverage between historic and modern data by randomly downsampling the number of reads included in bam files of the modern, higher-coverage samples using the DownsampleSam function in Picard tools (https://broadinstitute.github.io/picard/). Variants were called for historic and downsampled bam files using the same 'mpileup' and 'call' functions in bcftools as above.

VCF files of both the full data and downsampled dataset were then filtered in VCFtools to remove indels and retain only biallelic sites. We removed from the datasets: all SNPs with missing data, with a minimum quality score less than 20, exceeding the Hardy-Weinberg threshold of 0.01, a min mean depth <1, and a minimum depth < 5; individuals with mean coverage <4.5x; and individuals that appeared to be mis-identified migrants based on preliminary principal components analyses. The final dataset included 219 individuals and 196,151 SNPs. We applied identical filters to the downsampled dataset and generated a final downsampled dataset with 219 individuals and 130,317 SNPs.

Although we treated historic samples with the USER enzyme, sequencing errors could still be associated with certain sites in the genome. These include CpG sites or sites with C->T or G->A mutations (Briggs et al. 2009; van der Valk et al. 2019; Bi et al. 2019). We further filtered our datasets to remove all CpG SNPs from one dataset and removed all C->T and G->A mutations from another dataset in order to evaluate the influence of these potential biases on our results.

### SUPPLEMENTAL RESULTS:

*Summary of sequencing results:* Sequencing using our custom target array was highly successful for both historic and modern samples (Supplemental Table S1). In total, we generated 316.0 Gb (mean 4.27 Gb per individual) and 520.9 Gb (3.18 Gb per individual) of raw data for modern and historical samples, respectively. Across the four California subspecies of Savannah Sparrow, on average 73.0-73.7% of cleaned reads mapped to the target regions in modern samples and on average 70.4-77.4% of reads were on target in the historic samples (Supplemental Table S1). Sensitivity, the percent of target regions with at least one read mapping to it, was higher for modern samples (98.6-98.7%) than for historic samples (92.0-95.0%). Average coverage was about five times higher in modern samples (range: 40.5-73.7; mean: 55.0x) than historical samples (range: 2.0-29.4; mean: 11.0x). Downsampled modern individuals had an average coverage of 10x (Supplemental Figure S2). While there was no relationship between coverage and specificity (percent reads mapping to target regions), sensitivity did increase with greater coverage and plateaued above ~20x coverage (Supplemental Figure S1). For both historic and modern datasets the proportion of SNPs that were transitions were much higher than transversions. Certain transitions, such as C to T mutations, can be much more common in historical DNA due to deamination of Cs to Ts. Our data suggest that these mutation types were actually slightly more common in the modern dataset than historic dataset (Supplemental Figure S3). This was true both for the full dataset and data that was downsampled to achieve equal coverage between modern and historic datasets.

*Population structure:* For principal components analysis, restricting the data to just SNPs from exon regions, similar patterns of population structure remain regardless of whether it is a downsampled dataset, CpG sites are removed, or C->T and G->A mutations are removed (Supplemental Figure S4). Restricting the data to SNPs from nongenic target regions showed similar patterns of population structure along the first principal component with *P. s. beldingi,* Morro Bay, and the rest of California forming three distinct clusters. However, the second principal component instead separates modern from historical samples for each of the three clusters. This pattern is similar across all three non-genic datasets (Supplemental Figure S4). Greater structure between historic and modern samples in non-genic relative to exon regions could reflect increased drift in regions of the genome unconstrained by purifying selection.

SUPPLEMENTAL FIGURES

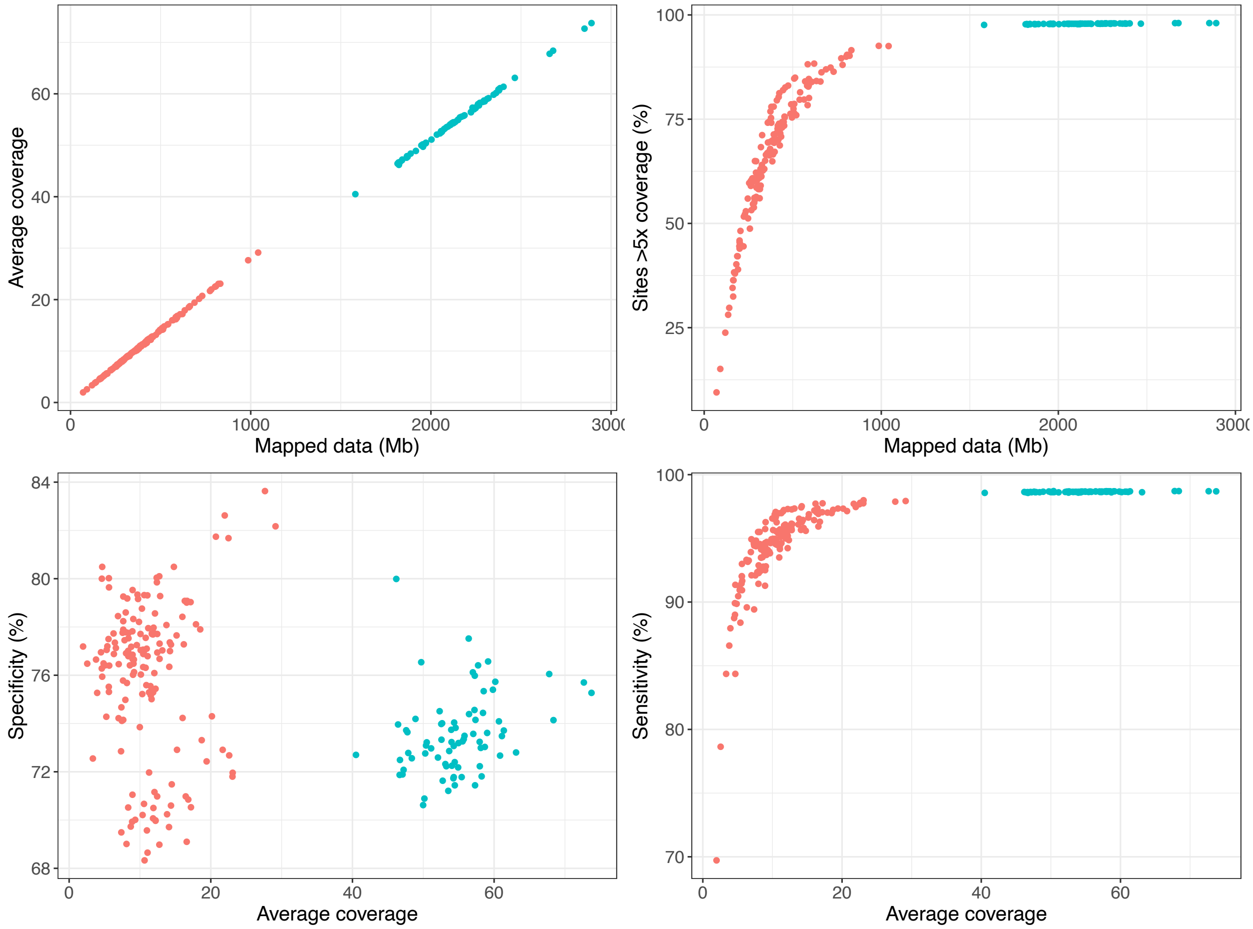


**Supplemental Figure S1:** Differences in sequencing success of target capture approach between modern samples extracted from fresh blood or tissue after the year 2000 (blue) and historic samples extracted from museum specimens before 1970 (red). Top left panel shows the relationship for average coverage and the amount of cleaned reads mapped to reference genome for each individual. Top right compares coverage at sites with >5x coverage and mapped data. Bottom left compares average coverage for each individual with specificity, the percent of reads mapping on-target. Bottom right compares average coverage with sensitivity, the percent of target regions with at least 1 read mapping.

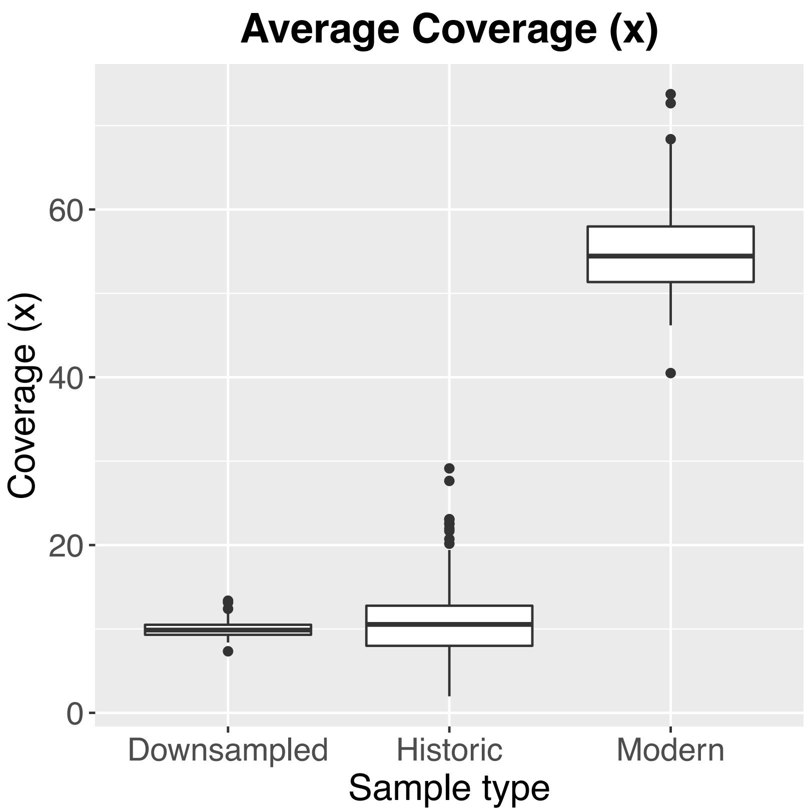


**Supplemental Figure S2:** Differences in coverage between modern samples extracted from fresh blood or tissue after the year 2000 and historic samples extracted from museum specimens before 1970. Downsampled refers to average coverage of modern samples after randomly downsampling the amount of mapped reads to approximate an equal amount of coverage to the historic samples.

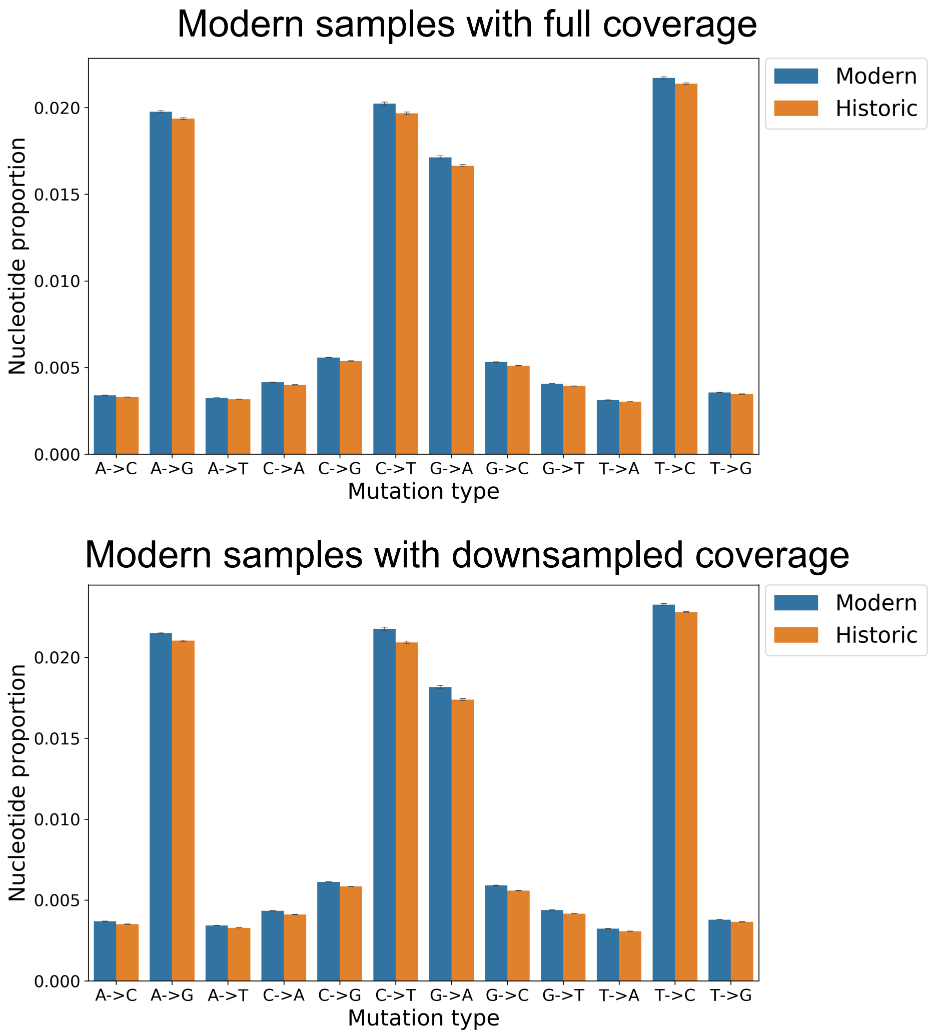


**Supplemental Figure S3:** Comparison of mutation patterns between modern (blue) and historic samples (orange). Bars correspond to each of the different mutation types with transitions representing a greater proportion of total SNPs than transversions. Top panel shows full dataset and lower panel shows mutation patterns following random downsampling of mapped reads.

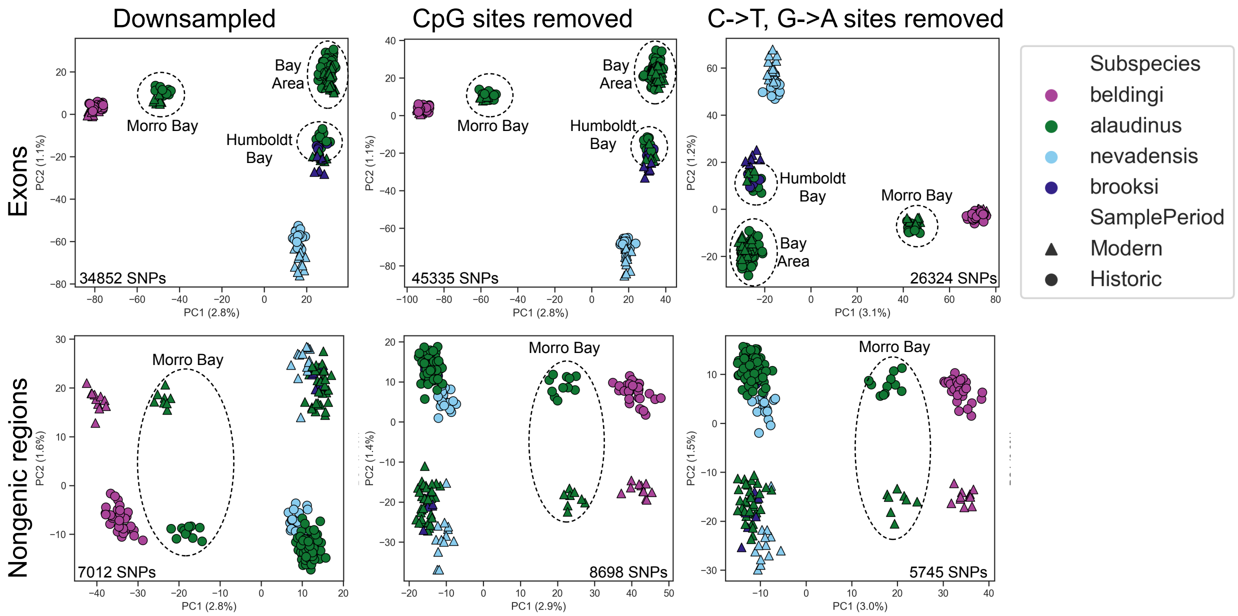


**Supplemental Figure S4:** Influence of different filtering regimes and kinds of data on patterns of population structure identified via principal components analysis. Top panels illustrate population structure within the exon dataset only and lower panels show patterns of structure based on analysis of the nongenic target regions. Downsampled dataset refers to the random downsampling of mapped reads in the modern dataset to ensure equal levels of coverage between sampling times. Middle panels show patterns of structure in datasets where all SNPs at CpG sites have been removed. Right most panels show results for datasets with all C->T and G->A SNPs removed. All datasets were pruned to remove SNPs within linkage of one another and final number of SNPs included in principal components analysis is listed on each figure. Colors correspond to different subspecies as illustrated in Fig. 1a. Dashed circles denote different clusters corresponding to different populations within *P. s. alaudinus.*

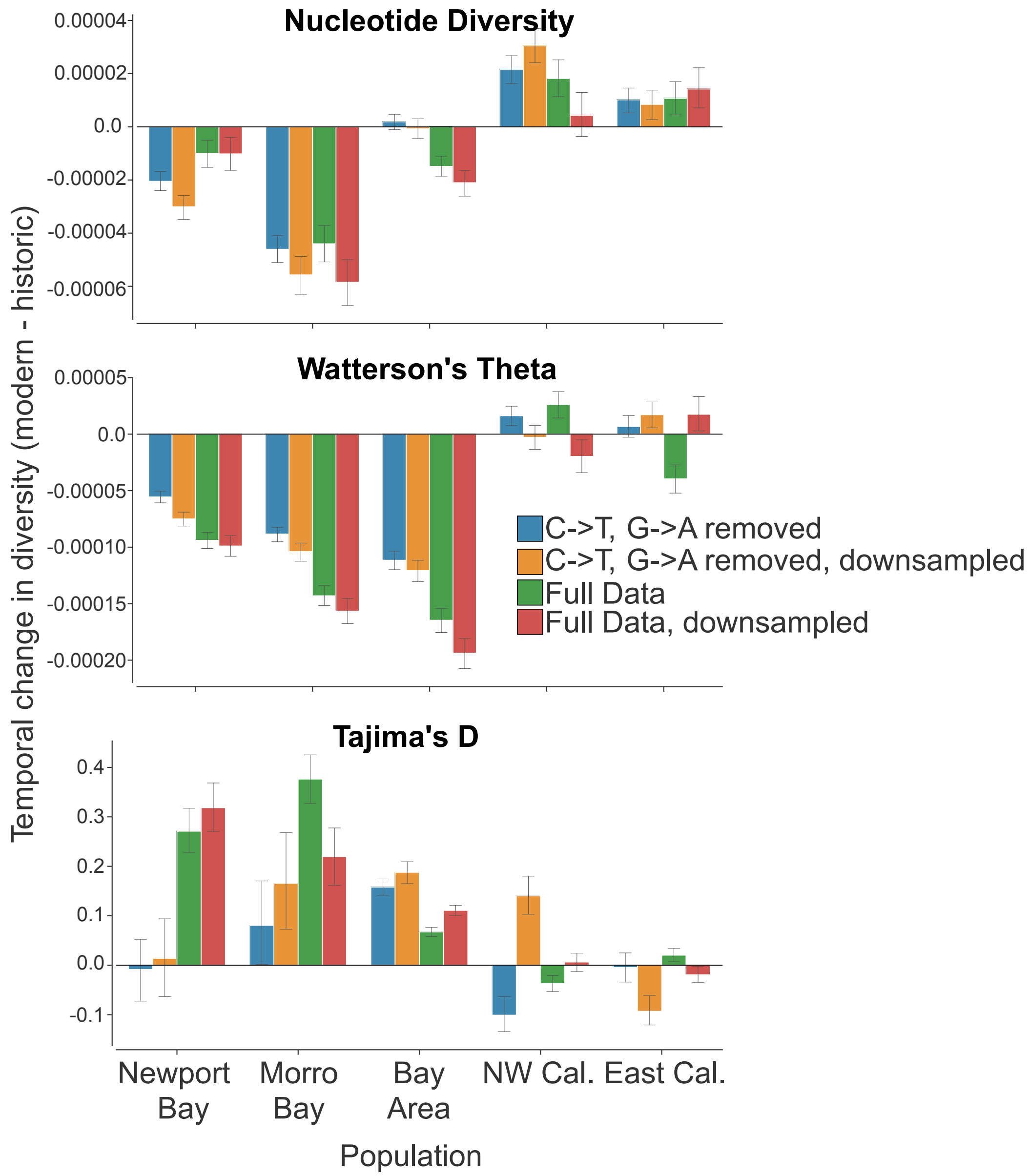


**Supplemental Figure S5:** Magnitude of temporal change in three metrics of genetic diversity across different populations and datasets with different filtering regimes. For nucleotide diversity, Watterson's theta, and Tajima's D the y-axis shows the difference between modern and historic estimates of each statistic. Populations on the x-axis correspond to the five population clusters identified in the PCA illustrated in Fig. 2b. Different colors of bars represent four different datasets that differed in whether coverage of modern samples was downsampled and/or if C->T or G->A mutations were removed.

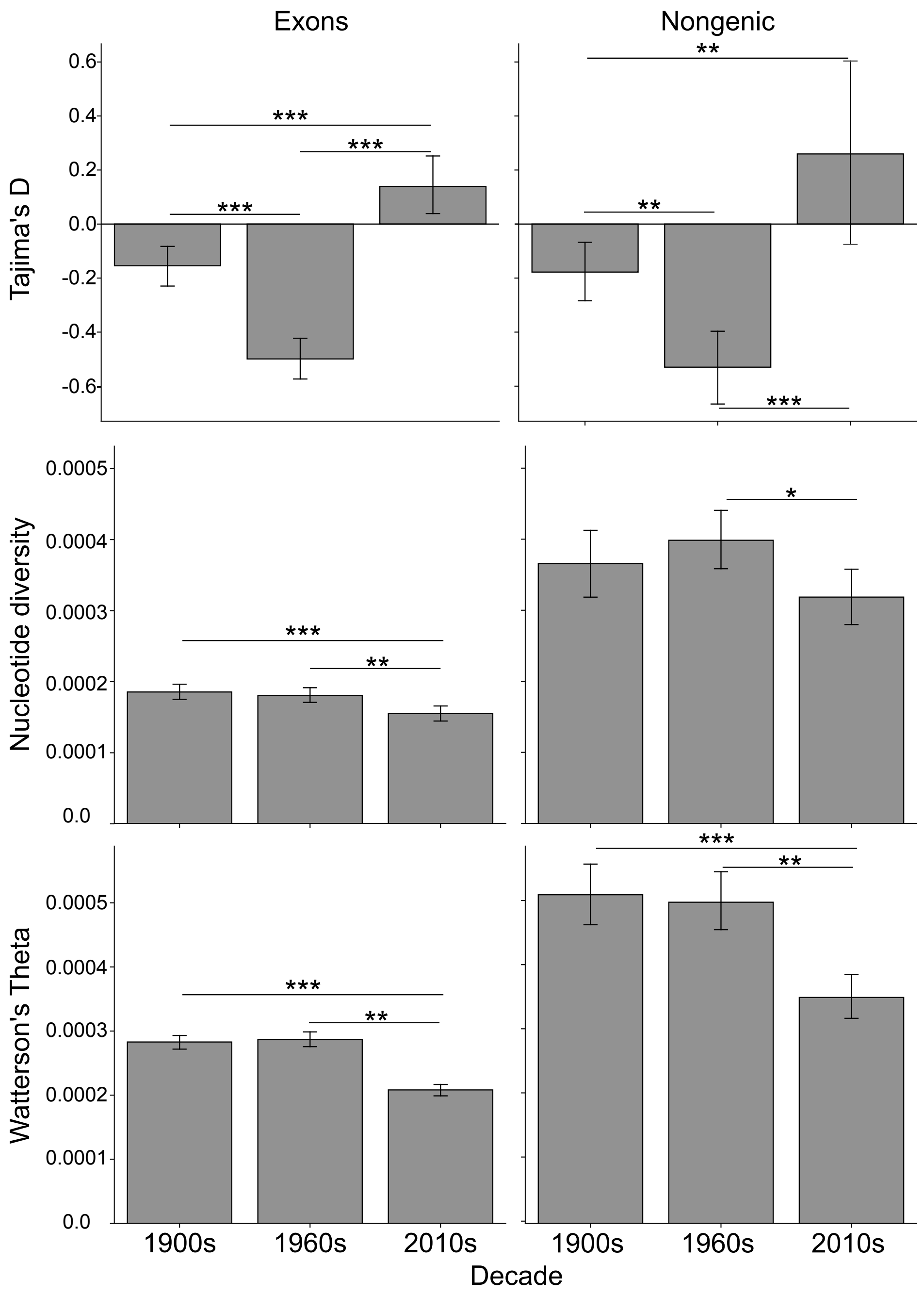


**Supplemental Figure S6:** Temporal variation in three metrics of genetic diversity in the Newport Bay population of Belding's Savannah sparrows (*P. s. beldingi*). Left panels show exon data and right panels nongenic data. Asterisks show significant differences between time periods based on ANOVA and TukeyHSD tests (*p<0.05; **p<0.01; ***: p<0.0001). Only significant differences are shown.

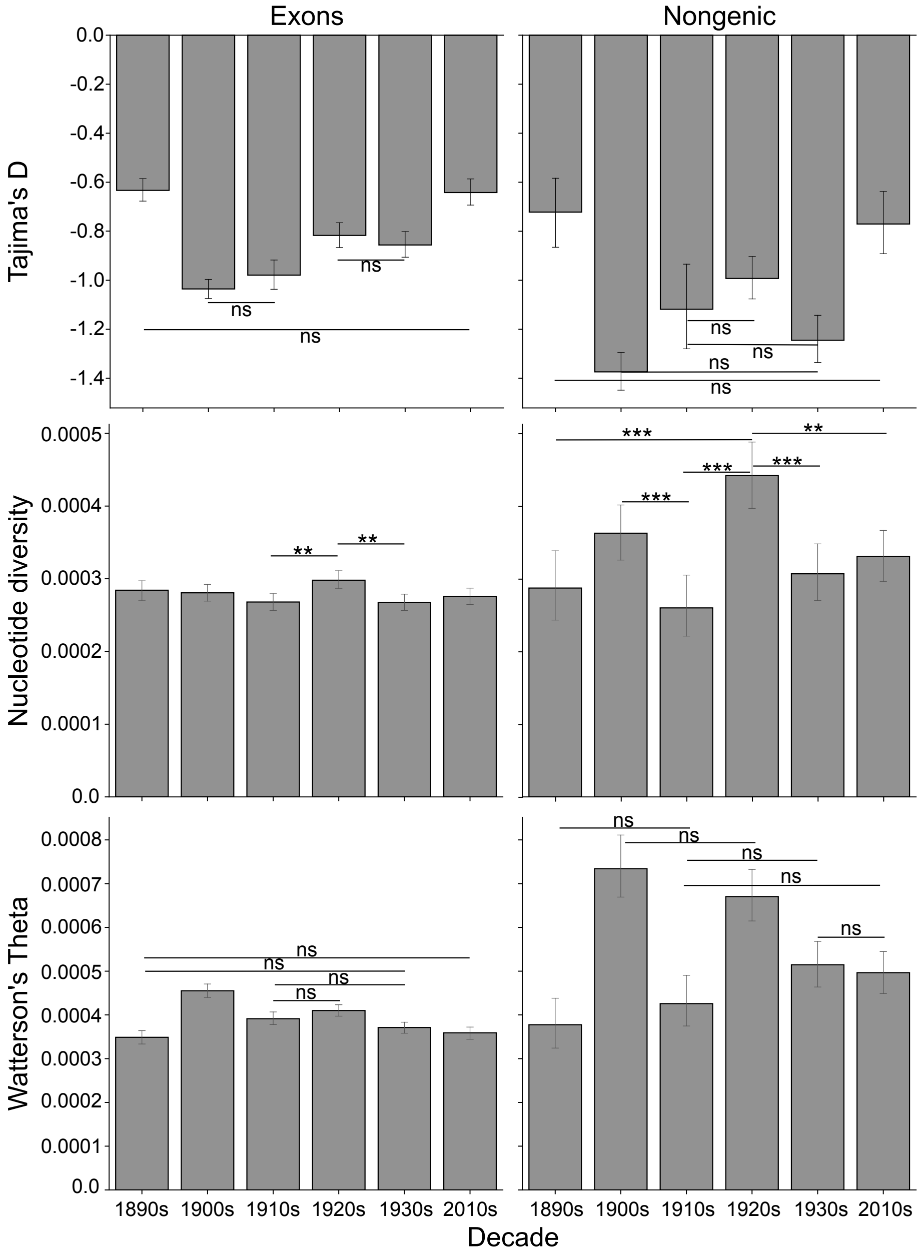


**Supplemental Figure S7:** Temporal variation in three metrics of genetic diversity in the San Francisco Bay population of Savannah sparrows (*P. s. alaudinus*). Left panels show exon data and right panels nongenic data. Asterisks show significant differences between time periods based on ANOVA and TukeyHSD tests (ns: non-significant; *p<0.05; **p<0.01; ***: p<0.0001). Only significant differences are shown for nucleotide diversity and only non-significant differences for Tajima's D and Watterson's Theta as most decades were statistically different from one another.

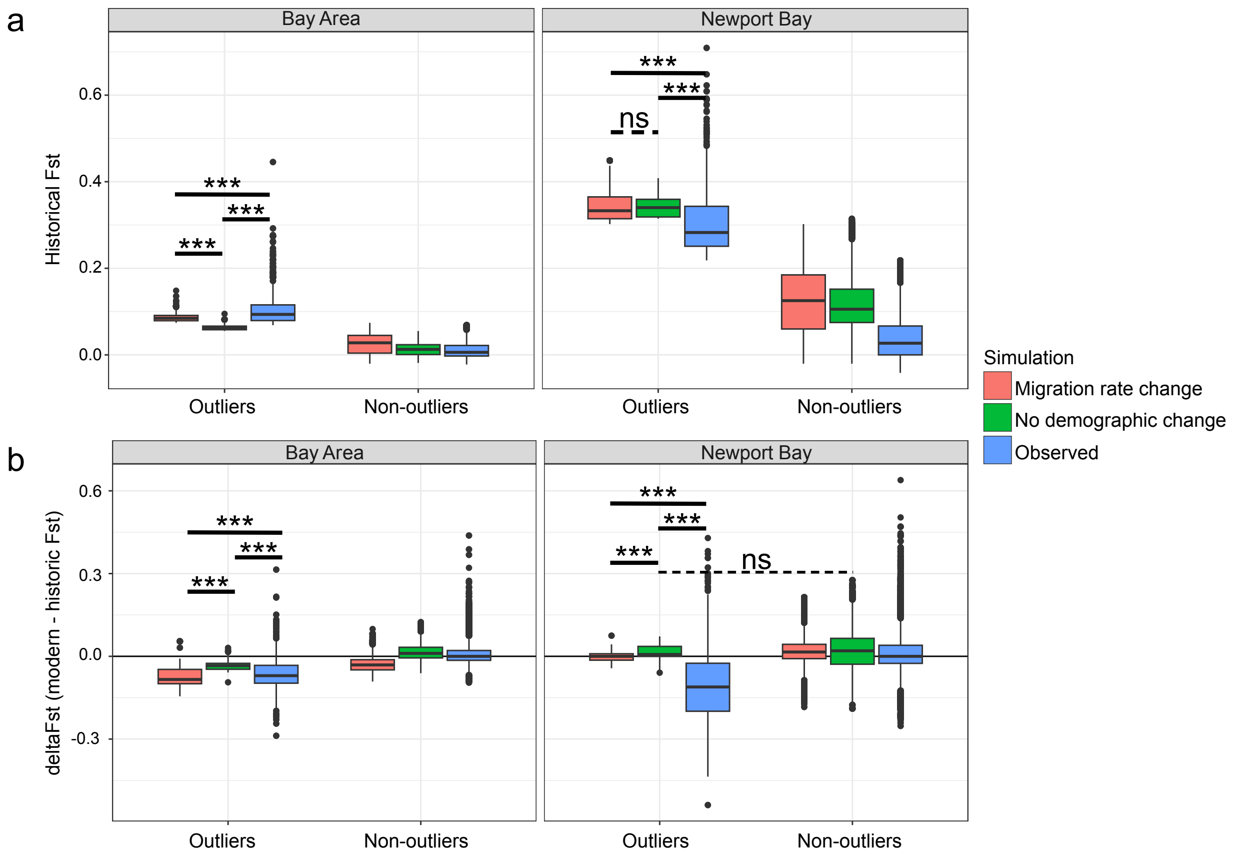


**Supplemental Figure S8:** Comparison of observed Fst divergence in the Bay Area and Newport Bay populations from the eastern California population and two simulated datasets. The first simulation models a neutral demographic history based on the historical best-fit demographic model with no demographic events occurring within the last 100 years. The second simulation models a migration rate change at 15 generations before the present with migration from eastern California to the Bay Area increasing to a rate of 0.15 (proportion of immigrants from east California in the Bay Area). Fst for historic and modern samples was estimated for all datasets in 5000bp sliding windows with a step size of 2500bp. (a) Patterns of historical Fst divergence with outliers (95 percentile) shown on left and non-outlier Fst windows on right in each plot. (b) Patterns of delta Fst (modern Fst - historic Fst) for outlier and non-outlier SNPs. For both the Bay Area and Newport Bay the observed negative change in delta Fst was greater than the expectation based on the neutral demographic model. For the Bay Area observed delta Fst remained slightly but significantly greater than the amount of change expected under the migration rate change model. Asterisks show significant differences between models based on ANOVA and TukeyHSD tests (ns: non-significant; *p<0.05; **p<0.01; ***: p<0.0001). Significant differences only shown for outlier loci.

SUPPLEMENTAL TABLES:

**Supplemental Table S1:** Summary table of sequencing metrics for California populations of the Savannah sparrow. Divided into modern samples extracted from fresh blood or tissue after the year 2000 and historic samples extracted from museum specimens before 1970. Raw data is the total amount of raw sequence data and mapped data is the total amount of cleaned data mapping to the *Zonotrichia albicollis* reference genome. Specificity refers to the percentage of reads that are mapping to target capture regions. Sensitivity is the percent of target regions with at least one read mapping to it. Average coverage is the mean coverage across all target regions from the custom capture array. All values are averages for each group and standard deviation is reported in parentheses.

**Modern**

| Subspecies | No. Samples | Raw data (Mb) | Mapped data (Mb) | Specificity (%) | Sensitivity (%) | Avg Cov. (X) |
|---|---|---|---|---|---|---|
| *P. s. alaudinus* | 44 | 4298.3 (522.4) | 2168.5 (263.9) | 73.7 (2.04) | 98.7 (0.03) | 55.4 (6.7) |
| *P. s. beldingi* | 11 | 4159.3 (376.1) | 2116.3 (178.8) | 73.2 (0.71) | 98.6 (0.03) | 54.1 (4.6) |
| *P. s. brooksi* | 6 | 4460.0 (242.4) | 2237.1 (110.3) | 73.0 (1.32) | 98.7 (0.01) | 57.2 (2.8) |
| *P. s. nevadensis* | 13 | 4180.5 (403.6) | 2076.3 (170.6) | 73.7 (0.58) | 98.7 (0.02) | 53.1 (4.3) |

**Historic**

| Subspecies | No. Samples | Raw data (Mb) | Mapped data (Mb) | Specificity (%) | Sensitivity (%) | Avg Cov. (X) |
|---|---|---|---|---|---|---|
| *P. s. alaudinus* | 103 | 3266.5 (1194.7) | 431.5 (179.0) | 76.2 (3.24) | 95.0 (2.84) | 12.1 (5.0) |
| *P. s. beldingi* | 33 | 3338.8 (1171.2) | 330.8 (127.9) | 75.1 (3.33) | 94.0 (2.72) | 9.3 (3.6) |
| *P. s. brooksi* | 6 | 3217.6 (596.8) | 382.4 (72.2) | 70.4 (0.78) | 94.4 (1.53) | 10.8 (2.0) |
| *P. s. nevadensis* | 19 | 2451.41 (785.5) | 297.3 (135.0) | 77.4 (1.51) | 92.0 (6.18) | 8.4 (3.8) |

**Supplemental Table S2:** The relationship between genetic diversity and amount of tidal marsh habitat. Comparison of Tajima's D (Tajd), nucleotide diversity (pi) and Watterson's theta (wattTheta) in relation to percent habitat lost for each estuary, historic habitat in log-transformed hectares (ha), and modern habitat in log-transformed hectares (ha).

| **delta diversity ~ % habitat loss** | | | |
|---|---|---|---|
| | $F_{(1,4)}$ | adj. $R^2$ | P-value |
| Tajd | 0.11 | -0.22 | 0.755 |
| pi | 3.91 | 0.37 | 0.12 |
| wattTheta | 0.04 | -0.23 | 0.84 |

| **delta diversity ~ historic habitat (log-ha)** | | | |
|---|---|---|---|
| | $F_{(1,4)}$ | adj. $R^2$ | P-value |
| Tajd | 0.006 | -0.25 | 0.93 |
| pi | 2.1 | 0.18 | 0.22 |
| wattTheta | 0.02 | -0.24 | 0.88 |

| **delta diversity ~ modern habitat (log-ha)** | | | |
|---|---|---|---|
| | $F_{(1,4)}$ | adj. $R^2$ | P-value |
| Tajd | 0.02 | -0.24 | 0.89 |
| pi | 0.82 | -0.04 | 0.42 |
| wattTheta | 0.00043 | -0.25 | 0.98 |

| **modern diversity ~ modern habitat (log-ha)** | | | |
|---|---|---|---|
| | $F_{(1,4)}$ | adj. $R^2$ | P-value |
| *Tajd* | *9.48* | *0.63* | *0.04* |
| pi | 2.7 | 0.25 | 0.17 |
| wattTheta | 7.14 | 0.55 | 0.056 |

| **modern diversity ~ historic habitat (log-ha)** | | | |
|---|---|---|---|
| | $F_{(1,4)}$ | adj. $R^2$ | P-value |
| *Tajd* | *11.29* | *0.67* | *0.028* |
| pi | 6.6 | 0.53 | 0.06 |
| *wattTheta* | *19.44* | *0.79* | *0.012* |

| **historic diversity ~ historic habitat (log-ha)** | | | |
|---|---|---|---|
| | $F_{(1,4)}$ | adj. $R^2$ | P-value |
| *Tajd* | *17.94* | *0.77* | *0.01* |
| pi | 2.75 | 0.26 | 0.17 |
| wattTheta | 7.33 | 0.56 | 0.053 |

**Supplemental Table S3**: Summary of genes found to be outliers in Fst, Dxy, and LFMM analyses. For each population detection of gene as an outlier in that population is marked by 1. Possible role of gene indicated if involved in kidney function (KF), kidney development (KD), or kidney injury & disease (KI).

| **Gene_name** | **Newport Bay** | **Morro Bay** | **Bay Area** | **Humboldt Bay** | **Function** | **Notes** |
|---|---|---|---|---|---|---|
| AKAP13 | | | | 1 | KI | Association with PKDisease |
| ATP6V0A1 | | | 1 | | KF | proximal tubule transporter, hypertension regulation |
| BRAF | 1 | 1 | | | KI | association with renal cancer |
| CCDC177 | 1 | | 1 | | | |
| CEBPG | | | 1 | | | |
| CNOT6L | | | 1 | | | |
| CPD | | 1 | | 1 | | |
| CRELD1 | 1 | | 1 | | | heart development |
| CTSA | | 1 | | | KF | conversion of angiotensin I->II, expressed in crab-eating frog |
| CUNH1orf226 | 1 | | | | | |
| DNAJB9 | | 1 | | | KI | Kidney injury & disease |
| ESR1 | 1 | 1 | 1 | | KF, KD | mediates AQP2 expression, kidney hypertrophy in knockout mice |
| FCGBP | | | 1 | 1 | KI | Kidney disease (macrophage) association |
| FMO5 | 1 | | | | | |
| FOXA1 | 1 | 1 | | | KI, KD, KF | kidney disease & injury, expressed in developing kidney, interacts with ESR1 in mammary duct development |
| FOXRED2 | 1 | | | | KI | Kidney disease & Injury |
| GMDS | 1 | | | | | |
| HAO2 | 1 | 1 | | | KI | kidney disease associations |
| HEG1 | | | | 1 | | |
| ICK | | 1 | | | KD? | Impacts SHH signaling |
| IL22RA2 | 1 | | 1 | | | |
| KLHL30 | 1 | | | | | |
| KNL1 | | | | 1 | | |
| LARP1 | | | 1 | | | interactions wtih mTORC1? |
| LOC102061812 (OR14A16-like) | 1 | | | | | |
| LOC102063133 (CSNK1A1) | | | 1 | 1 | KI | kidney disease association |
| LOC102064430 (APOBEC1) | | | 1 | | | |
| LOC102064557 | 1 | | | | | |

| | | | | | | |
|---|---|---|---|---|---|---|
| LOC102065258 (ZNF271) | 1 | | | | | |
| LOC102065621 (CCNT1) | | 1 | | | | |
| LOC102067227 | | | 1 | 1 | | |
| LOC102068722 (CXCL8) | 1 | | | | KI | response to kidney injury |
| LOC102071624 (TWIST1) | 1 | | | | KI, KD | kidney injury & disease, renal & craniofacial development |
| LOC102072224 | | | 1 | | | |
| LOC102072901 (MFHAS1) | 1 | | | | KI | alleviates inflammation and renal fibrosis |
| LOC102073186 (Thymic CPV3) | 1 | | | | | |
| LOC102074067 (MFSD6) | | | 1 | | | |
| **LOC102074318 (FCGBP-like)** | 1 | | 1 | 1 | | |
| LOC102075054 (CHST9) | | 1 | | | | |
| LOC106629462 (ELL2) | | | 1 | 1 | | |
| LOC106629890 | 1 | | | | | |
| MPL | | | 1 | | | |
| MRPL3 | 1 | | | | | |
| MYO9A | | | 1 | | KF | regulator of kidney tubule function |
| NDST4 | 1 | 1 | | | | Colon epithelium development; hepearin biosynthesis |
| NGLY1 | | 1 | | | KF | Aquaporin regulation, hypoosmotic stress response |
| NYAP2 | 1 | | | | | |
| PCNT | | 1 | | | KI | kidney disease |
| PDE12 | 1 | | | | | |
| PLD5 | | 1 | | 1 | | |
| PLEKHO2 | | 1 | | | | |
| RAD50 | | | | 1 | KF | activated in response to hyperosmotic cellular stress, involved in complex that repairs double-stranded DNA breaks |
| RASSF3 | | 1 | | | | |
| RCC1 | 1 | 1 | | | KI | association with renal disease |
| RD3L | | | 1 | | | |
| RFC4 | 1 | 1 | | | | |
| SENP1 | 1 | | | | KI | association with kidney injury |
| SMO | 1 | | | | KD | Kidney Development |
| SOX10 | 1 | 1 | | | | Neural crest development, interacts with ESR1,TWIST, etc. |

| | | | | | | |
|---|---|---|---|---|---|---|
| SPARCL1 | 1 | | | | KI | expressed in response to kidney injury |
| SPEG | | | | 1 | KI | GWAS with GFR |
| TAF3 | 1 | | | | | |
| TCF20 | | 1 | | | | |
| TIMM23B | 1 | | 1 | | KI | |
| TMEM106C | | | 1 | | | |
| TMEM159 | 1 | 1 | | | | |
| TMEM198 | | 1 | | | | |
| TSC1 | | 1 | | | KI | Involved in PKD and other renal diseases |
| TTN | 1 | | | | | pulmonary hypertension |
| VASH1 | | | 1 | | KI | Kidney protection |

**Supplemental Table S4**: Historical outlier statistics for Newport Bay, Orange Co. (*P. s. beldingi*).

| Gene name | LFMM p-value | Fst | ΔFst | Dxy | π salt | π fresh | Scaffold | Start |
|---|---|---|---|---|---|---|---|---|
| BRAF | 0.0013 | 0.362 | -0.093 | 4.10E-04 | 1.24E-04 | 3.99E-04 | NW_005081546.1 | 11158492 |
| CCDC177 | 0.0001 | 0.227 | -0.123 | 4.61E-04 | 1.57E-04 | 5.56E-04 | NW_005081537.1 | 13984146 |
| CCDC177 | 0.0001 | 0.266 | -0.221 | 2.78E-04 | 5.79E-05 | 3.50E-04 | NW_005081537.1 | 13986646 |
| CRELD1 | 0.0014 | 0.530 | -0.217 | 2.72E-04 | 7.79E-05 | 1.78E-04 | NW_005081600.1 | 678483 |
| CUNH1orf226 | 0.0033 | 0.229 | -0.154 | 4.00E-04 | 2.75E-04 | 3.42E-04 | NW_005081730.1 | 632775 |
| ESR1 | 0.0332 | 0.512 | -0.007 | 2.91E-04 | 3.79E-05 | 2.46E-04 | NW_005081596.1 | 1906681 |
| ESR1 | 0.0332 | 0.512 | -0.007 | 2.91E-04 | 3.79E-05 | 2.46E-04 | NW_005081596.1 | 1909181 |
| FMO5 | 0.0468 | 0.343 | -0.321 | 1.11E-03 | 1.14E-03 | 3.15E-04 | NW_005083203.1 | 2821 |
| FOXA1 | 0.0272 | 0.259 | -0.033 | 3.91E-04 | 1.97E-04 | 3.83E-04 | NW_005081537.1 | 4264146 |
| FOXA1 | 0.0272 | 0.262 | -0.032 | 3.85E-04 | 1.97E-04 | 3.71E-04 | NW_005081537.1 | 4266646 |
| FOXRED2 | 0.0190 | 0.518 | -0.050 | 2.89E-04 | 1.67E-04 | 1.11E-04 | NW_005081603.1 | 492681 |
| GMDS | 0.0464 | 0.254 | -0.111 | 4.69E-04 | 3.83E-04 | 3.18E-04 | NW_005081542.1 | 5786791 |
| GMDS | 0.0464 | 0.254 | -0.111 | 4.69E-04 | 3.83E-04 | 3.18E-04 | NW_005081542.1 | 5789291 |
| HAO2 | 0.0384 | 0.413 | -0.247 | 3.22E-04 | 1.65E-04 | 2.13E-04 | NW_005081570.1 | 1022849 |
| HAO2 | 0.0384 | 0.445 | -0.245 | 2.92E-04 | 1.12E-04 | 2.13E-04 | NW_005081570.1 | 1025349 |
| IL22RA2 | 0.0007 | 0.255 | -0.190 | 3.05E-04 | 1.32E-04 | 3.23E-04 | NW_005081611.1 | 2700496 |
| KLHL30 | 0.0334 | 0.315 | 0.073 | 2.94E-04 | 2.44E-04 | 1.59E-04 | NW_005081617.1 | 1527669 |
| LOC102061812 | 0.0233 | 0.251 | -0.169 | 2.80E-04 | 1.94E-04 | 2.26E-04 | NW_005082406.1 | 1686 |
| LOC102064557 | 0.0114 | 0.266 | -0.002 | 3.07E-03 | 2.52E-03 | 1.99E-03 | NW_005082576.1 | 22642 |
| LOC102065258 | 0.0230 | 0.264 | 0.032 | 6.94E-03 | 7.00E-03 | 3.22E-03 | NW_005083076.1 | 13260 |
| LOC102068537 | 0.0240 | 0.334 | 0.105 | 2.79E-04 | 2.75E-04 | 9.66E-05 | NW_005081808.1 | 238924 |
| LOC102071624 | 0.0433 | 0.253 | -0.104 | 2.77E-04 | 1.37E-04 | 2.77E-04 | NW_005083478.1 | 19628 |
| LOC102072901 | 0.0103 | 0.222 | -0.030 | 4.58E-04 | 1.34E-04 | 5.78E-04 | NW_005081665.1 | 932088 |
| LOC102073186 | 0.0484 | 0.340 | -0.181 | 2.72E-04 | 1.73E-04 | 1.86E-04 | NW_005081585.1 | 634061 |
| LOC102073186 | 0.0484 | 0.340 | -0.181 | 2.72E-04 | 1.73E-04 | 1.86E-04 | NW_005081585.1 | 636561 |
| LOC102074318 | 1.99E-06 | 0.271 | -0.004 | 5.81E-04 | 1.25E-04 | 7.22E-04 | NW_005082169.1 | 24479 |
| LOC102074318 | 0.0005 | 0.276 | 0.007 | 3.35E-04 | 1.05E-04 | 3.80E-04 | NW_005082169.1 | 26979 |
| LOC106629890 | 0.0339 | 0.282 | -0.177 | 9.59E-04 | 5.70E-04 | 8.08E-04 | NW_005081804.1 | 587286 |
| MRPL3 | 0.0288 | 0.312 | -0.200 | 2.79E-04 | 1.83E-04 | 2.01E-04 | NW_005081540.1 | 17932957 |
| MRPL4 | 0.0288 | 0.312 | -0.200 | 2.79E-04 | 1.83E-04 | 2.01E-04 | NW_005081540.1 | 17935457 |
| NDST4 | 4.09E-05 | 0.293 | 0.166 | 5.33E-04 | 6.51E-04 | 1.03E-04 | NW_005081789.1 | 191766 |
| NDST4 | 4.09E-05 | 0.293 | 0.166 | 5.33E-04 | 6.51E-04 | 1.03E-04 | NW_005081789.1 | 194266 |
| NYAP2 | 0.007 | 0.269 | -0.062 | 2.79E-04 | 2.47E-04 | 1.61E-04 | NW_005081614.1 | 993618 |
| NYAP2 | 0.007 | 0.245 | -0.043 | 3.09E-04 | 2.47E-04 | 2.19E-04 | NW_005081614.1 | 996118 |
| PDE12 | 0.044 | 0.283 | -0.013 | 2.91E-04 | 1.46E-04 | 2.71E-04 | NW_005081564.1 | 2435547 |
| RCC1 | 0.013 | 0.256 | -0.190 | 4.55E-04 | 3.05E-04 | 3.73E-04 | NW_005081926.1 | 47814 |

| | | | | | | | | |
|---|---|---|---|---|---|---|---|---|
| RCC1 | 0.013 | 0.355 | -0.269 | 2.93E-04 | 1.69E-04 | 2.08E-04 | NW_005081926.1 | 50314 |
| RFC4 | 0.001 | 0.358 | -0.080 | 3.27E-04 | 1.71E-04 | 2.50E-04 | NW_005081559.1 | 4835513 |
| RFC4 | 0.001 | 0.358 | -0.080 | 3.27E-04 | 1.71E-04 | 2.50E-04 | NW_005081559.1 | 4838013 |
| SENP1 | 0.019 | 0.272 | -0.156 | 2.92E-04 | 1.34E-04 | 2.91E-04 | NW_005082306.1 | 11868 |
| SMO | 0.042 | 0.242 | -0.043 | 2.11E-03 | 1.61E-03 | 1.59E-03 | NW_005083929.1 | 192 |
| SOX10 | 0.018 | 0.225 | 0.295 | 3.81E-04 | 2.95E-04 | 2.96E-04 | NW_005081603.1 | 1210181 |
| SPARCL1 | 0.016 | 0.280 | -0.159 | 2.89E-04 | 1.72E-04 | 2.45E-04 | NW_005081565.1 | 6259796 |
| SPARCL1 | 0.016 | 0.292 | -0.167 | 2.78E-04 | 1.72E-04 | 2.22E-04 | NW_005081565.1 | 6262296 |
| TAF3 | 0.005 | 0.278 | -0.271 | 3.30E-04 | 4.00E-05 | 4.37E-04 | NW_005081772.1 | 497051 |
| TIMM23B | 0.010 | 0.301 | -0.269 | 3.07E-04 | 6.00E-05 | 3.69E-04 | NW_005081541.1 | 18638896 |
| TIMM23B | 0.010 | 0.319 | -0.284 | 2.89E-04 | 6.00E-05 | 3.34E-04 | NW_005081541.1 | 18641396 |
| TMEM159 | 0.029 | 0.227 | 0.036 | 3.06E-04 | 2.28E-04 | 2.45E-04 | NW_005081585.1 | 131561 |
| TMEM159 | 0.029 | 0.218 | 0.039 | 3.18E-04 | 2.28E-04 | 2.68E-04 | NW_005081585.1 | 134061 |
| TTN | 0.038 | 0.246 | -0.128 | 4.72E-04 | 1.55E-04 | 5.57E-04 | NW_005081554.1 | 5627881 |

**Supplemental Table S5**: Historical outlier statistics for Morro Bay, San Luis Obispo Co. (*P. s. beldingi/alaudinus*).

| Gene name | LFMM p-value | Fst | ΔFst | Dxy | π salt | π fresh | Scaffold | Start |
|---|---|---|---|---|---|---|---|---|
| BRAF | 0.001 | 0.227 | -0.029 | 4.01E-04 | 3.99E-04 | 2.20E-04 | NW_005081546.1 | 11158492 |
| CPD | 0.028 | 0.177 | 0.070 | 4.72E-04 | 5.92E-04 | 1.84E-04 | NW_005081728.1 | 226877 |
| CTSA | 0.042 | 0.190 | -0.057 | 3.20E-04 | 3.06E-04 | 2.13E-04 | NW_005081656.1 | 1617675 |
| CTSA | 0.042 | 0.190 | -0.057 | 3.20E-04 | 3.06E-04 | 2.13E-04 | NW_005081656.1 | 1620175 |
| DNAJB9 | 8.95E-05 | 0.250 | -0.145 | 2.95E-04 | 1.80E-04 | 2.63E-04 | NW_005081539.1 | 164698 |
| ESR1 | 0.033 | 0.201 | 0.307 | 2.84E-04 | 2.46E-04 | 2.09E-04 | NW_005081596.1 | 1906681 |
| ESR1 | 0.033 | 0.201 | 0.307 | 2.84E-04 | 2.46E-04 | 2.09E-04 | NW_005081596.1 | 1909181 |
| FOXA1 | 0.027 | 0.233 | -0.067 | 3.07E-04 | 3.83E-04 | 8.74E-05 | NW_005081537.1 | 4264146 |
| FOXA1 | 0.027 | 0.238 | -0.067 | 3.01E-04 | 3.71E-04 | 8.74E-05 | NW_005081537.1 | 4266646 |
| HAO2 | 0.038 | 0.166 | -0.009 | 2.87E-04 | 2.13E-04 | 2.65E-04 | NW_005081570.1 | 1022849 |
| ICK | 0.032 | 0.189 | -0.163 | 3.91E-04 | 3.77E-04 | 2.58E-04 | NW_005081569.1 | 6349679 |
| LOC102065621 | 0.003 | 0.159 | -0.163 | 8.58E-03 | 6.09E-03 | 8.34E-03 | NW_005082267.1 | 49954 |
| LOC102075054 | 0.015 | 0.239 | -0.189 | 3.27E-04 | 1.93E-04 | 3.05E-04 | NW_005081537.1 | 17514146 |
| NDST4 | 4.09E-05 | 0.166 | 0.155 | 3.78E-04 | 1.03E-04 | 5.26E-04 | NW_005081789.1 | 191766 |
| NDST4 | 4.09E-05 | 0.166 | 0.155 | 3.78E-04 | 1.03E-04 | 5.26E-04 | NW_005081789.1 | 194266 |
| NGLY1 | 0.020 | 0.243 | -0.206 | 4.63E-04 | 2.12E-04 | 4.88E-04 | NW_005081540.1 | 13847957 |
| NGLY1 | 0.020 | 0.243 | -0.206 | 4.63E-04 | 2.12E-04 | 4.88E-04 | NW_005081540.1 | 13850457 |
| PCNT | 0.002 | 0.283 | -0.132 | 3.02E-04 | 2.66E-04 | 1.67E-04 | NW_005081578.1 | 3158670 |
| PLD5 | 0.042 | 0.172 | -0.168 | 3.16E-04 | 3.15E-04 | 2.08E-04 | NW_005081620.1 | 1320998 |
| PLD5 | 0.042 | 0.167 | -0.158 | 3.25E-04 | 3.15E-04 | 2.26E-04 | NW_005081620.1 | 1323498 |
| PLEKHO2 | 0.018 | 0.185 | 0.034 | 3.95E-04 | 2.00E-04 | 4.44E-04 | NW_005081550.1 | 9436025 |
| RASSF3 | 0.049 | 0.159 | -0.015 | 3.26E-04 | 3.38E-04 | 2.10E-04 | NW_005081539.1 | 5359698 |
| RCC1 | 0.013 | 0.225 | -0.147 | 3.22E-04 | 2.08E-04 | 2.91E-04 | NW_005081926.1 | 50314 |
| RFC4 | 0.001 | 0.251 | -0.158 | 2.86E-04 | 2.50E-04 | 1.78E-04 | NW_005081559.1 | 4835513 |
| RFC4 | 0.001 | 0.251 | -0.158 | 2.86E-04 | 2.50E-04 | 1.78E-04 | NW_005081559.1 | 4838013 |
| SOX10 | 0.018 | 0.323 | 0.132 | 4.11E-04 | 2.96E-04 | 2.61E-04 | NW_005081603.1 | 1210181 |
| TCF20 | 0.014 | 0.168 | 0.091 | 3.15E-04 | 3.31E-04 | 1.93E-04 | NW_005081603.1 | 3177681 |
| TMEM159 | 0.029 | 0.258 | -0.125 | 3.22E-04 | 2.45E-04 | 2.33E-04 | NW_005081585.1 | 131561 |
| TMEM159 | 0.029 | 0.243 | -0.115 | 3.43E-04 | 2.68E-04 | 2.51E-04 | NW_005081585.1 | 134061 |
| TMEM198 | 0.036 | 0.201 | 0.031 | 2.89E-04 | 2.27E-04 | 2.35E-04 | NW_005081562.1 | 6420712 |
| TSC1 | 0.031 | 0.195 | -0.098 | 3.42E-04 | 2.96E-04 | 2.55E-04 | NW_005081757.1 | 409066 |
| TSC1 | 0.031 | 0.199 | -0.100 | 3.36E-04 | 2.84E-04 | 2.55E-04 | NW_005081757.1 | 411566 |

**Supplemental Table S6**: Historical outlier statistics for San Francisco Bay Area (*P. s. alaudinus*).

| Gene name | LFMM p-value | Fst | ΔFst | Dxy | π salt | π fresh | Scaffold | Start |
|---|---|---|---|---|---|---|---|---|
| ATP6V0A1 | 0.012 | 0.085 | -0.094 | 3.04E-04 | 2.27E-04 | 3.30E-04 | NW_005081766.1 | 618487 |
| ATP6V0A1 | 0.012 | 0.088 | -0.098 | 2.90E-04 | 2.21E-04 | 3.07E-04 | NW_005081766.1 | 620987 |
| CCDC177 | 8.78E-05 | 0.095 | -0.049 | 3.31E-04 | 2.50E-04 | 3.50E-04 | NW_005081537.1 | 13986646 |
| CEBPG | 0.038 | 0.069 | -0.064 | 3.88E-04 | 3.75E-04 | 3.48E-04 | NW_005081693.1 | 961001 |
| CNOT6L | 0.001 | 0.100 | -0.076 | 2.99E-04 | 2.64E-04 | 2.73E-04 | NW_005081808.1 | 248924 |
| CNOT6L | 0.001 | 0.100 | -0.077 | 2.99E-04 | 2.64E-04 | 2.73E-04 | NW_005081808.1 | 251424 |
| CRELD1 | 0.001 | 0.152 | -0.120 | 3.21E-04 | 3.79E-04 | 1.66E-04 | NW_005081600.1 | 675983 |
| CRELD1 | 0.001 | 0.149 | -0.117 | 3.27E-04 | 3.79E-04 | 1.78E-04 | NW_005081600.1 | 678483 |
| ESR1 | 0.033 | 0.200 | -0.057 | 2.98E-04 | 2.31E-04 | 2.46E-04 | NW_005081596.1 | 1906681 |
| ESR1 | 0.033 | 0.200 | -0.057 | 2.98E-04 | 2.31E-04 | 2.46E-04 | NW_005081596.1 | 1909181 |
| FCGBP | 0.000 | 0.081 | -0.005 | 3.45E-04 | 2.78E-04 | 3.56E-04 | NW_005081730.1 | 1050275 |
| IL22RA2 | 0.001 | 0.147 | -0.023 | 2.90E-04 | 1.70E-04 | 3.23E-04 | NW_005081611.1 | 2700496 |
| LARP1 | 0.004 | 0.086 | -0.092 | 2.87E-04 | 2.88E-04 | 2.36E-04 | NW_005081551.1 | 8926162 |
| LOC102063133 | 0.025 | 0.071 | -0.066 | 3.75E-04 | 2.67E-04 | 4.30E-04 | NW_005081603.1 | 1337681 |
| LOC102064430 | 0.048 | 0.078 | -0.023 | 3.52E-04 | 3.19E-04 | 3.30E-04 | NW_005081713.1 | 170744 |
| LOC102064430 | 0.048 | 0.078 | -0.023 | 3.52E-04 | 3.19E-04 | 3.30E-04 | NW_005081713.1 | 173244 |
| LOC102067227 | 0.044 | 0.080 | -0.077 | 1.89E-02 | 1.68E-02 | 1.80E-02 | NW_005082889.1 | 22450 |
| LOC102072224 | 0.001 | 0.077 | -0.057 | 4.07E-03 | 2.22E-03 | 5.28E-03 | NW_005084020.1 | 456 |
| LOC102074067 | 0.014 | 0.084 | -0.064 | 2.88E-04 | 2.61E-04 | 2.67E-04 | NW_005081702.1 | 782 |
| LOC102074318 | 1.99E-06 | 0.107 | -0.035 | 6.65E-04 | 4.66E-04 | 7.22E-04 | NW_005082169.1 | 24479 |
| LOC102074318 | 0.001 | 0.099 | 0.003 | 2.94E-04 | 1.50E-04 | 3.80E-04 | NW_005082169.1 | 26979 |
| LOC106629462 | 0.004 | 0.125 | -0.099 | 3.16E-04 | 1.75E-04 | 3.78E-04 | NW_005081601.1 | 4188644 |
| LOC106629462 | 0.002 | 0.160 | -0.036 | 4.69E-04 | 2.25E-04 | 5.64E-04 | NW_005081601.1 | 4191144 |
| MPL | 0.018 | 0.073 | -0.067 | 4.44E-04 | 1.86E-04 | 6.37E-04 | NW_005081544.1 | 8395220 |
| MPL | 0.018 | 0.078 | -0.074 | 4.20E-04 | 1.60E-04 | 6.14E-04 | NW_005081544.1 | 8397720 |
| MYO9A | 0.002 | 0.125 | -0.071 | 3.63E-04 | 2.64E-04 | 3.71E-04 | NW_005081634.1 | 2252704 |
| MYO9A | 0.002 | 0.119 | -0.071 | 3.79E-04 | 2.85E-04 | 3.83E-04 | NW_005081634.1 | 2255204 |
| RD3L | 0.011 | 0.072 | -0.066 | 3.04E-04 | 2.52E-04 | 3.12E-04 | NW_005081549.1 | 10026571 |
| TIMM23B | 0.010 | 0.069 | -0.082 | 3.57E-04 | 3.31E-04 | 3.34E-04 | NW_005081541.1 | 18641396 |
| TMEM106C | 0.001 | 0.115 | -0.126 | 2.81E-04 | 3.18E-04 | 1.80E-04 | NW_005082297.1 | 19180 |
| VASH1 | 0.030 | 0.071 | -0.042 | 3.35E-04 | 2.45E-04 | 3.77E-04 | NW_005081537.1 | 2439146 |
| VASH1 | 0.030 | 0.079 | -0.047 | 2.95E-04 | 2.14E-04 | 3.30E-04 | NW_005081537.1 | 2441646 |

**Supplemental Table S7**: Historical outlier statistics for Humboldt Bay, Humboldt Co. (*P. s. alaudinus*).

| Gene name | LFMM p-value | Fst | ΔFst | Dxy | π salt | π fresh | Scaffold | Start |
|---|---|---|---|---|---|---|---|---|
| AKAP13 | 0.022 | 0.129 | -0.179 | 3.36E-04 | 2.96E-04 | 2.91E-04 | NW_005081550.1 | 5586025 |
| CPD | 0.028 | 0.096 | -0.005 | 5.34E-04 | 5.92E-04 | 3.74E-04 | NW_005081728.1 | 226877 |
| FCGBP | 0.000 | 0.116 | -0.116 | 3.70E-04 | 3.56E-04 | 2.98E-04 | NW_005081730.1 | 1050275 |
| HEG1 | 0.041 | 0.150 | -0.039 | 3.32E-04 | 3.25E-04 | 2.39E-04 | NW_005081842.1 | 280128 |
| KNL1 | 0.048 | 0.098 | -0.082 | 3.09E-04 | 2.57E-04 | 3.01E-04 | NW_005081787.1 | 392415 |
| LOC102063133 | 0.025 | 0.108 | -0.086 | 3.85E-04 | 4.30E-04 | 2.57E-04 | NW_005081603.1 | 1337681 |
| LOC102063133 | 0.025 | 0.100 | -0.080 | 4.15E-04 | 4.30E-04 | 3.17E-04 | NW_005081603.1 | 1340181 |
| LOC102067227 | 0.044 | 0.161 | -0.154 | 1.90E-02 | 1.80E-02 | 1.39E-02 | NW_005082889.1 | 22450 |
| LOC102074318 | 1.99E-06 | 0.149 | 0.007 | 6.61E-04 | 7.22E-04 | 4.04E-04 | NW_005082169.1 | 24479 |
| LOC106629462 | 0.002 | 0.104 | -0.021 | 4.94E-04 | 5.64E-04 | 3.22E-04 | NW_005081601.1 | 4191144 |
| PLD5 | 0.042 | 0.096 | -0.113 | 3.51E-04 | 3.15E-04 | 3.19E-04 | NW_005081620.1 | 1320998 |
| PLD5 | 0.042 | 0.094 | -0.111 | 3.61E-04 | 3.15E-04 | 3.39E-04 | NW_005081620.1 | 1323498 |
| RAD50 | 0.020 | 0.115 | -0.139 | 3.39E-04 | 2.86E-04 | 3.15E-04 | NW_005081551.1 | 3338662 |
| SPEG | 0.037 | 0.129 | -0.116 | 5.73E-04 | 3.51E-04 | 6.47E-04 | NW_005081562.1 | 6310712 |
| SPEG | 0.037 | 0.133 | -0.119 | 4.85E-04 | 3.16E-04 | 5.26E-04 | NW_005081562.1 | 6313212 |
